# H3.3 deposition counteracts the replication-dependent enrichment of H3.1 at chromocenters in embryonic stem cells

**DOI:** 10.1101/2024.07.04.601905

**Authors:** S. Arfè, T. Karagyozova, A. Forest, H. Hmidan, E. Meshorer, J.-P. Quivy, G. Almouzni

**Affiliations:** Institut Curie, PSL University, Sorbonne Université, CNRS, Nuclear Dynamics, 75005 Paris, France; Department of Genetics, The Alexander Silberman Institute for Life Science, and the Edmond and Lily Safra Center for Brain Sciences (ELSC), The Hebrew University of Jerusalem, Jerusalem, Israel 9190400; Center for Neural Science and Medicine, Department of Biomedical Sciences, Cedars-Sinai Medical Center, Los Angeles, CA 90048, USA; Physiology and pharmacology department, college of medicine, Al-Quds University, Jerusalem, Palestine

## Abstract

Chromocenters in mouse cells are membrane-less nuclear compartments that represent typical heterochromatin stably maintained during the cell cycle. Here, we explore how histone H3 variants, replicative H3.1/H3.2 or replacement H3.3, mark these domains during the cell cycle. In mouse embryonic stem cells (ESCs), neuronal precursor cells (NPCs) as well as immortalized 3T3 cells, we find a strong and distinct H3.1 enrichment at chromocenters, with some variation in ESCs. Mechanistically, this H3.1 selective enrichment depends on the DNA Synthesis Coupled (DSC) deposition pathway operating in S phase. Yet, this selective enrichment is challenged when we target H3.3 deposition through the DNA Synthesis Independent (DSI) deposition pathway mediated by HIRA. Altering the H3.1/H3.3 equilibrium at chromocenters in ESCs affects its heterochromatin properties leading to mitotic defects. We thus reveal opposing mechanisms for H3.1 and H3.3 deposition with different enforcement according to cell cycle and potency which determine their ratio at chromocenters and are critical for genome stability and cell survival.

## Introduction

Chromatin states and their plasticity is associated with distinct cell fates ^1^. During mammalian development, starting from a fertilized egg, a series of cellular divisions, along with the acquisition of distinct specialized functions, provide the variety of cell types necessary to form a complete adult organism. Each cell displays distinct chromatin features using a versatile basic unit, the nucleosome. Indeed, the nucleosomal core particle, with 147 bp of genomic DNA wrapped around a histone octamer made of an H3-H4 tetramer flanked by two H2A-H2B dimers ^2^, can provide different flavors with the choice of histone variants and their histone post-translation modifications (PTMs) ^3,4^. In mammals, two major non-centromeric histone H3 (H3.1/H3.2 and H3.3) variants, display distinct distribution patterns across the genome and use distinct deposition pathways. Notably, replicative variants, such as H3.1/H3.2, are deposited via a DNA Synthesis Coupled (DSC) pathway promoted by the chaperone CAF-1 complex mainly during S phase and show a broad genome-wide distribution ^5^. In contrast, H3.3, a replacement variant, is incorporated in a DNA Synthesis-Independent (DSI) fashion either by the chaperone HIRA at active chromatin regions and specialized nuclear domains ^5,6^ or by the DAXX-ATRX complex at constitutive heterochromatin regions, including telomeres, retrotransposons, and pericentric heterochromatin ^7–10^. Heterochromatin is typically associated with high levels of constitutive heterochromatin marks, including H3K9me3 and HP1 proteins which disruption is often associated with increased DNA damage and defects in chromosome segregation ^11,12^. In addition, heterochromatin integrity is also linked to chromatin replication since the perturbation of the replicative CAF-1 histone chaperone disrupts chromocenter formation in both ESCs and mouse embryos ^13^.

Chromocenters are membrane-less nuclear domains visible as DAPI-stained foci in interphase nuclei. In mouse cells, they correspond to the clustering of pericentric domains from different chromosomes forming typical constitutive heterochromatin ^14^. The pericentric domains, mostly composed of major satellite DNA repeats, flank the most centric region which contains minor satellite DNA repeats and is enriched in CENP-A, the centromeric variant. These chromosomal landmarks play essential roles in genome organization and stability ^14,15^. During development and cellular differentiation, chromocenters undergo dynamic remodeling, reflecting changes in chromatin and nuclear architecture ^16–22^. The importance of histone variants in cell fate choices during development and disease ^23^ has further raised the importance to consider their relative distribution and its regulation. Indeed, the differential deposition of histone variants and distinct PTMs has emerged as an important player in establishing euchromatin and heterochromatin during development ^16^. For example, facultative heterochromatin marks, such as H3K27me3, are critical for maintaining lineage-specific gene expression programs and H3.3 has been implicated in marking these regions ^24,25^. While H3.3 exhibits a distinct genomic enrichment pattern, H3.1/H3.2 variants are more broadly distributed and associated dynamically in the chromatin of totipotent and pluripotent cells ^26,27^. Interestingly, in plants, chromocenters show a specific enrichment in H3.1 ^28^. In mouse embryonic stem cells (ESCs), during replication in heterochromatin regions, H3.1/H3.2 recycling and maintenance is ensured both before and after differentiation, while active regions do not show such a stable maintenance ^29,30^. Thus, understanding the mechanisms determining histone variant dynamics at distinct subnuclear regions such as chromocenters can shed light onto means to achieve distinct genome marking in different cells.

In this study, we investigated the nuclear distribution of H3.1 and H3.3 variants, focusing on their relative enrichment at chromocenters throughout the cell cycle and in different cellular states. Leveraging microscopy and chromatin immunoprecipitation followed by sequencing (ChIP-seq) analyses, we monitored the spatial and temporal dynamics of histone variant in pluripotent mouse ESCs, neuronal precursor cells (NPCs), and immortalized differentiated fibroblast cells (NIH-3T3). While we detected both histone variants in the whole nucleus, only the replicative variants H3.1/H3.2 systematically stood out at chromocenters, regardless of the cell potency state. Mechanistically, we provide evidence that DSC deposition is the major driver for the replicative H3 variant accumulation at chromocenters while excluding H3.3. Moreover, by forcing H3.3 deposition via the HIRA deposition pathway we challenged this pattern. This H3 variant unbalance led to heterochromatin alterations triggering mitotic defects in mouse ESCs. We discuss how competition between histone variant deposition can ensure proper chromatin organization with a significant impact on cell function.

## Results

### Mouse chromocenters show a specific enrichment in replicative H3 variant

To explore the subnuclear localization of the H3.1 and H3.3 variants in cells with different differentiation potentials, we introduced SNAP-HA-tagged H3.1 and H3.3 under the control of a Tet-ON system in mouse ESCs (Fig. S1a). We first verified that the Doxycycline-induced expression of exogenous H3.1-SNAP-HA and H3.3-SNAP-HA did not interfere with the pluripotency potential of these engineered ESCs as attested by the Oct3/4 detection (Fig. S1b). We could readily visualize the nuclear H3.1-SNAP-HA and H3.3-SNAP-HA using SNAP-pulse labeling *in vivo* with TMR (Fig. S1c) 18 h after the addition of doxycycline to the ESCs with no further increase in fluorescence signal beyond 48h of treatment (Fig. S1c), indicating that the amounts of exogenously expressed H3.1 and H3.3 had reached equilibrium after 48 hours of doxycycline treatment (Fig. S1c). Western blot analysis using total cell extracts further showed that, under these conditions, exogenous H3.1 or H3.3 histones represented less than 5% of the total histone pool (Fig. S1d) minimally interfering with the endogenous pool. We next monitored the incorporation of these exogenous H3.1 and H3.3 into chromatin by pre-extracting soluble histones before fixation ^31,32^ (Fig 1a, b). By imaging, we observed two main patterns for H3.1: a marked enrichment at chromocenters (‘Enriched’ pattern) or diffuse distribution throughout the nucleus (‘Even’) (Fig. 1b, upper panels 1 and 2). In contrast, H3.3 rather showed either a weak depletion at chromocenters (‘Excluded’) or a homogenously diffuse distribution throughout the nucleus (‘Even’) (Fig. 1b, lower panels 3 and 4). Linescan quantification confirmed these patterns for the nuclear distribution of exogenous H3 variants (Fig. S1e). Importantly, using specific H3.1/H3.2 and H3.3 antibodies in non-engineered SNAP cells (parental cells), we reproduced the same patterns for endogenous H3.1/H3.2 and H3.3 (Fig. 1c). We next examined the nuclear localization of the two H3 variants after differentiation. We differentiated ESCs into neural progenitor cells (NPCs) and generated NIH-3T3 cell lines constitutively expressing SNAP-HA-tagged H3.1 and H3.3. The loss of Oct3/4 detection indicated a loss of pluripotency of NPCs (Fig. S1b) and the identical expression levels of exogenous H3 variants in NPCs and 3T3 cells compared to ESCs (Fig. S1d). The localization of exogenous H3 variants in NPCs and 3T3 cells proved identical to ESCs, with a clear enrichment at chromocenters for H3.1 (along with HP1α) in contrast to H3.3 (Fig. S2a). Furthermore, we confirmed this subnuclear localization for endogenous H3.1/H3.2 and H3.3 variants in all three cell types, ESCs, NPCs, and NIH-3T3 (Fig. 1c). We further quantified at chromocenters the immunofluorescence signal corresponding to the detection of endogenous H3.1/H3.2 (and not H3.3) in pluripotent (ESCs) and differentiated cells (3T3). For this, we considered the ratio between signal intensity at chromocenters and the rest of the nucleus using a 3D-FIED (3-dimension-fluorescence intensity enrichment at domains) method ^33^. The value for H3.1/H3.2 signal ratio at chromocenters consistently reached a value above 1, indicative of enrichment with a slightly lower value in ESCs when compared to 3T3 cells (Avg ratio 1.42 +/-0.28 and 1.59 +/-0.30, respectively) (Fig. S2b, left panel). In contrast, we found a ratio for H3.3 rather around 1 indicating no particular enrichment or even a weak depletion (1.01 +/-0.08 for 3T3 and 0.94 +/-0.04 for ESCs) (Fig. S2b, right panel). Since we noticed some heterogeneity and cells did not display solely ‘enriched’ (for H3.1/H3.2) or ‘excluded’ (for H3.3) but had also an ‘even’ pattern (Fig. 1a,b), we quantified the proportion of cells with ‘enriched’, ‘even’, and ‘excluded’ patterns in ESCs, NPCs, and 3T3 cells. We performed the analysis both on endogenous variants using H3.1/H3.2 and H3.3 specific antibodies in cells non-expressing exogenous histones (Fig. 1a, Fig. S1e) and exogenous SNAP-HA tagged H3 variants by TMR pulse labeling *in vivo* (Fig. 1c, Fig. S2a). We found that for endogenous H3.3, ∼75% of cells displayed ‘even’ and ∼25% ‘exclusion’ patterns at PHC for every cell line analyzed (ES, NPC, and 3T3) (Fig. 1d left panel). We obtained a similar distribution for exogenous H3.3 (Fig. 1d right panel). For H3.1, every cell line showed ‘enriched’ and ‘even’ patterns, with virtually no ‘excluded’ pattern (Fig. 1d). However, in contrast to H3.3, H3.1 showed different proportions of each pattern in the three cell lines. While the differentiated cell lines (NPCs and 3T3) had ∼85% of ‘enriched’ and ∼15% of ‘even’ patterns, the pluripotent ESC lines, instead, displayed reproducibly a lower frequency of cells with “enriched” patterns (∼70%). Importantly we found this difference between differentiated cells and ESCs for both endogenous (Fig. 1d, left panel) and exogenously expressed H3.1 (Fig. 1d, right panel). These results are consistent with the increased enrichment of H3.1/H3.2 at chromocenters in differentiated 3T3 cells when compared to pluripotent ESCs (Fig. S2b). They further reveal a less prominent H3.1/H3.2 enrichment at chromocenters in ESCs compared to differentiated cells, possibly counterbalanced by H3.3 to ensure H3 variant density.

**Figure 1.**
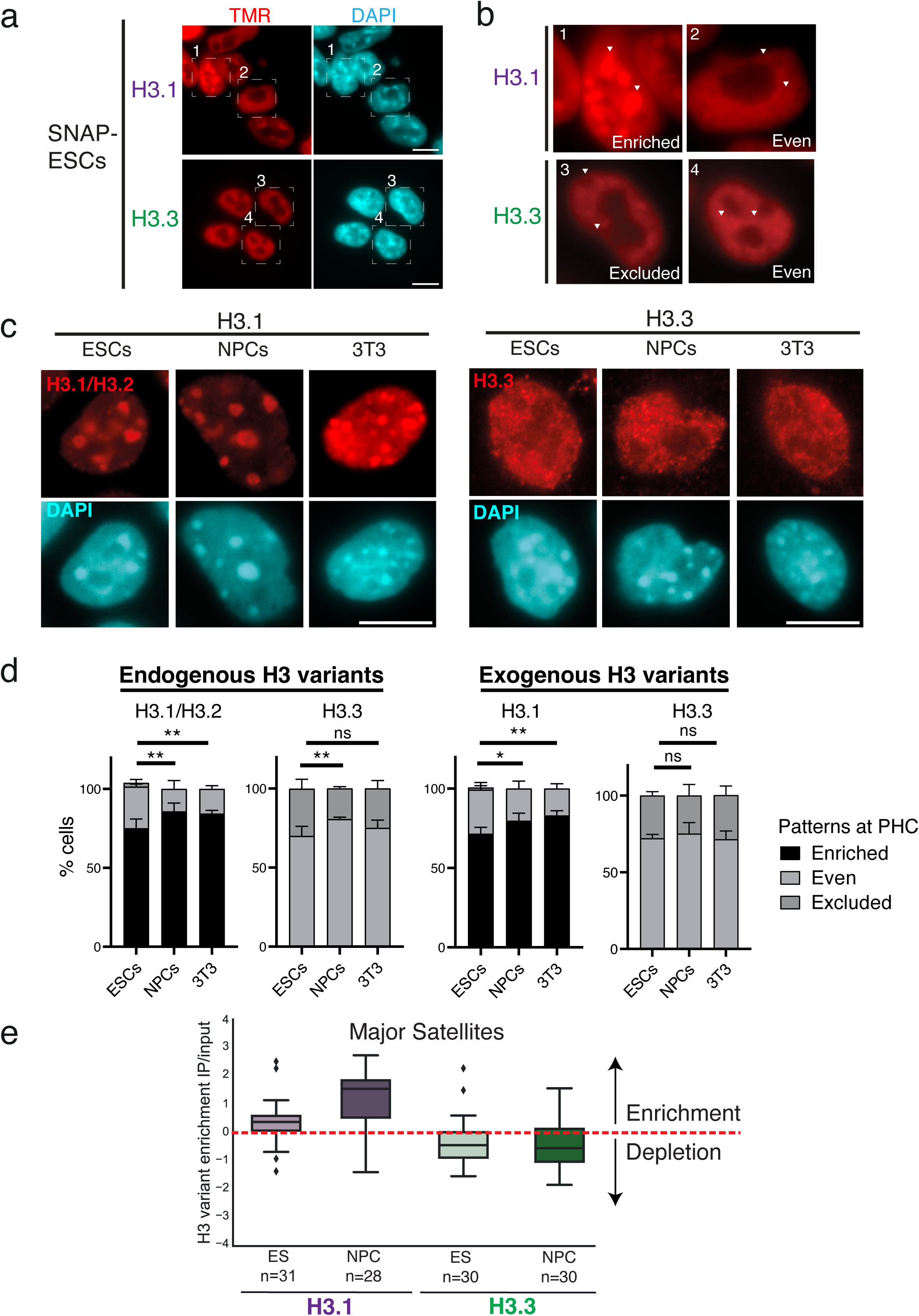
Histone variant H3.1 is enriched at chromocenters in mouse cells while H3.3 is not. (**a**) Representative wide-field epifluorescence images of Pulse-labeled (TMR) H3.1 and H3.3-SNAP (red) and DNA counterstaining with DAPI (cyan) in mouse ESCs. Single planes are shown, and squared boxes indicate nuclei on the right panels. Scale bar: 10 µm. (**b**) Insets of representative ESC nuclei showing typical H3 patterns at chromocenters for H3.1/H3.3. Definition of the three observed patterns at chromocenters, based on IF signal intensity in euchromatin and at chromocenters: 1) Enriched: strong H3 signal at chromocenters compared to euchromatin, 2) and 4): Even: Equal H3 intensity signal at chromocenters and euchromatin, 3) Excluded: Weaker H3 signal intensity at chromocenters compared to euchromatin. White arrows point to clusters of pericentric chromatin (or chromocenters) previously identified as DAPI-dense regions. Scale bar: 10 µm. See also supplementary material Fig. S1 and S2 for characterization of the cell lines and definition of observed patterns. (**c**) Representative epifluorescent images of endogenous H3.1/H3.2 (left) and H3.3 (right) detected with antibodies (red) in ESCs, neuronal progenitor cells (NPCs) derived from ESCs, and NIH-3T3 cells, followed by DNA counterstaining (DAPI, cyan). Scale bar: 10 µm. (**d**) Quantitative analysis of the percentage of cells exhibiting recurrent patterns of endogenous and exogenous H3 variants at PHC in different cell backgrounds. Data show the mean and standard deviations (s.d.) from at least 3 biologically independent experiments; >100 nuclei were quantified per condition. ANOVA tests were used for statistical analysis: ns (p > 0.05), * (p < 0.05), ** (p < 0.01). Scale bar: 10 mm. D (**e**) Quantification of H3 variants at Major Satellite repeat elements throughout differentiation. The Y-axis displays the H3 enrichment represented as a Z-score of log2 enrichment of IP over input. n represents the number of repeat elements at each differentiation step. The color palette from light to dark pattern reflects the differentiation conditions analyzed.

Next in a genome-wide approach, we exploited the SNAP-HA-H3.1 and SNAP-HA-H3.3 tagged ESC lines and performed SNAP capture followed by high-throughput sequencing (SNAP capture-seq) ^34,35^ to map H3.1 and H3.3 associated DNA sequences (Fig. S2c). We analyzed and quantified the distribution of H3.1 and H3.3 at the major satellite DNA repeats for both ESCs and NPCs. Importantly, we observed a steady increase of H3.1 enrichment during differentiation, with relatively weak depletion of H3.3 at chromocenters (Fig. 1e). Thus, the genomic data confirm a specific H3.1 enrichment over chromocenter-associated repetitive elements that are maintained from pluripotent to differentiated cells. We conclude that irrespective of the amount of the H3 variant (endogenous or exogenous), the cell differentiation status, or the presence of a genetic tag, the specific subnuclear localization of histone H3 variants remains a robust parameter. Importantly, this analysis also underlines a dynamic enrichment at chromocenters during differentiation, where a reciprocal behavior between H3.1 and H3.3 is observed.

### H3.1 enrichment at chromocenters varies during the cell cycle in ESCs contrasting with the steady distribution in differentiated cells

The fact that, in an asynchronous cell population, we did not detect in every cell a strong H3.1 enrichment at chromocenters suggested possible cell cycle variations. Indeed, cell cycle dynamics in mESCs is very different from NPCs and fibroblasts ∼12 hr cycle with very short G1 ^36^. Thus, we monitored H3 patterns in parallel with cell cycle markers. We used Aurora B and EdU labeling, to discriminate the G1 (Aurora B-/EdU-), S (Aurora B-/EdU+), and G2 cells (Aurora B+/EdU-) (Fig. 2a). The spatiotemporal organization of DNA replication foci (based on EdU pattern) further enables to follow S phase progression with (i) “Early S-phase” showing a high density of small foci throughout the nucleus; (ii) “Mid S-phase” with typical ring-shaped staining around the chromocenters; and (ii) “Late S-phase” with a few large foci at the nuclear periphery ^37–39^. Additionally, within the S-phase cell population, we could distinguish cells with chromocenters that had not yet replicated (‘Early’) from those undergoing replication (‘Mid’) and/or had already experienced replication (‘Late’). This analysis showed that H3.1 nuclear patterns changed with cell cycle progression (Fig. 2b, left panels) while H3.3 was stable (Fig. 2b, right panel). We estimated the proportion of cells displaying ‘enriched’, ‘even’, and ‘excluded’ patterns at chromocenters for H3.1 and found that NPCs and 3T3 cells presented a high proportion (>80%) of ‘enriched’ patterns at all cell cycle stages (G1, Mid S-phase, Late S-phase, and G2) except for Early S-phase (∼50% ‘enriched’) (Fig. 2c, top panels). Surprisingly in ESCs, >80% of cells in Mid-S, Late-S, and G2 phases displayed an ‘enriched’ pattern, like NPCs and 3T3 cells, while the proportion of cells with an ‘enriched’ pattern decreased to 50% in G1. This is in sharp contrast with non-pluripotent NPCs and 3T3 cells which maintain the ‘enriched’ pattern in G1 (∼ 95%) (Fig. 2c). In addition, ESCs showed the lowest proportion of H3.1 ‘enriched’ pattern (∼10%) during early S-phase amongst all cell lines analyzed. When monitoring H3.3, a reciprocal picture emerged, namely cells with the highest H3.1 ‘enriched’ pattern correlated with cells showing the largest fraction of the H3.3 ‘excluded’ pattern (Fig. 2c, H3.3 bottom panels). Conversely, when we observed the lowest H3.1 ‘enriched’ pattern in G1 and Early S, most cells did not exhibit H3.3 ‘exclusion’. (Fig. 2c, H3.3 bottom panels).

**Figure 2.**
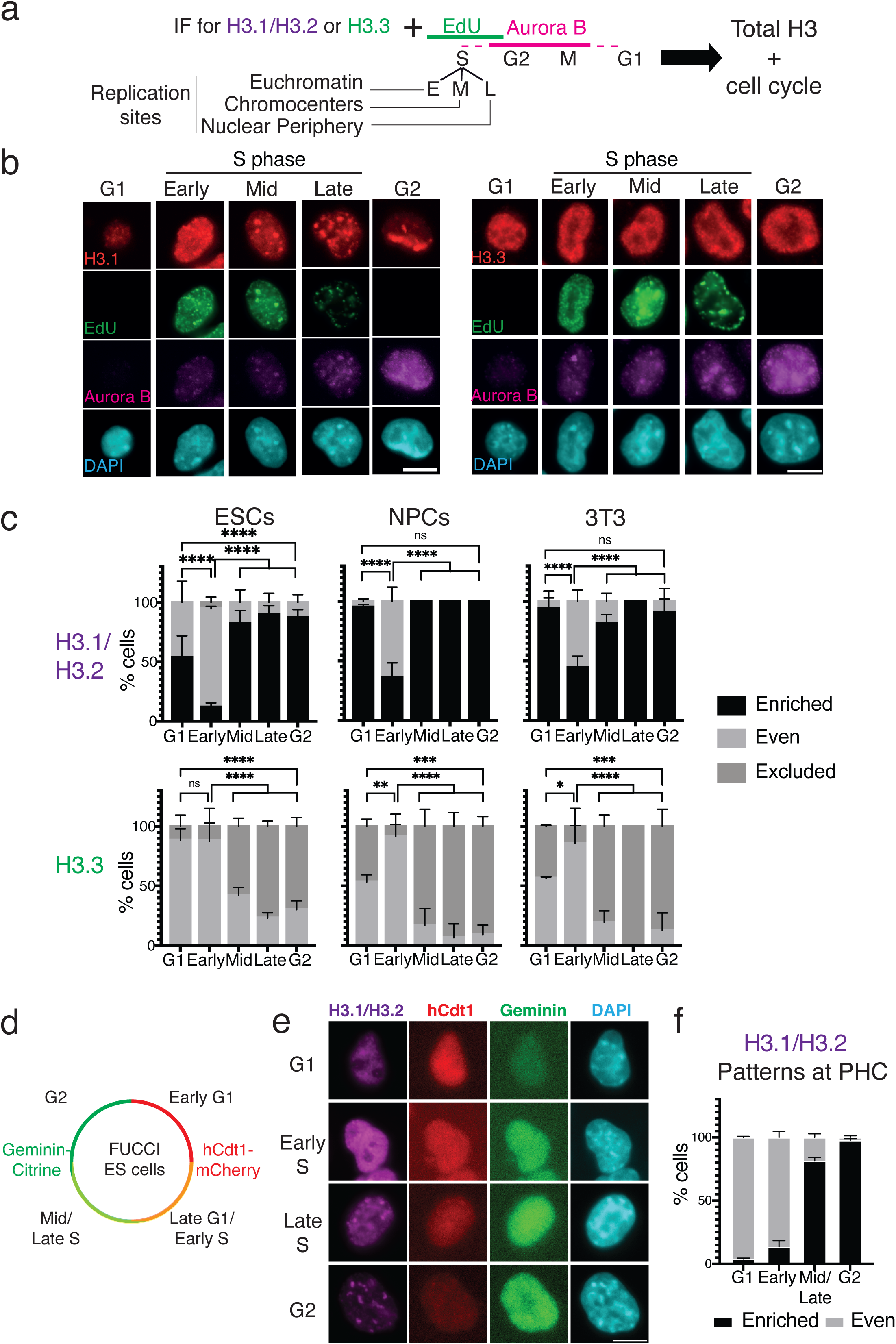
H3.1 enrichment at chromocenters follows chromocenter replication status. (**a**) Experimental scheme for visualizing endogenous H3.1/H3.2 or H3.3 during the S phase. Aurora B and EdU staining allowed to resolve the cell cycle stage of individual cells. Cells are scored as G1 (negative to Aurora B and EdU), S phase (EdU positive), or G2 (negative to EdU and positive to Aurora B). S-phase designations are based on EdU S phase patterns with the Early S phase (E) defined by a diffused staining (euchromatin); Mid S phase (M) by a focused EdU labeling around DAPI foci (chromocenters); and Late S phase (L) by specific foci staining at the nuclear periphery. (**b**) Left: Representative immunofluorescence images of ESCs after *in vivo* labeling with EdU (green), immunofluorescence staining of H3.1/H3.2 and H3.3 (red), Aurora B (magenta), and DNA (DAPI, cyan). Scale bars, 10 μm. (**c**) Quantitative analysis of the proportion of cells displaying H3 variant enrichment patterns at PHC in different cell lines (ESCs, NPCs, NIH-3T3) during the G1, S, and G2 phases. (**d**) Schematic of FUCCI ESCs allowing visualization by microscopy of hCdt1-mCherry and Geminin-Citrine during the cell cycle. Cells are scored as G1 (mCherry++/Citrine-), Late G1/Early S (mCherry+/Citrine+), Mid/Late S (mCherry-/Citrine+), or G2 (mCherry-/Citrine++). (**e**) Representative immunofluoresce images of FUCCI ESCs expressing hCdt1 (red) or Geminin (green) with immunofluorescent staining of H3.1/H3.2 (magenta) in G1, Early S, Late S, and G2 phases. Scale bars, 10 μm. (**f**) Quantitative analysis of the percentage of FUCCI ESCs exhibiting recurrent H3.1/H3.2 patterns at chromocenters throughout the cell cycle. All data show the mean and SD of at least 3 biologically independent experiments in >100 nuclei quantified per condition. ANOVA tests were used for statistical analysis: ns (p > 0.05), * (p < 0.05), *** (p < 0.001), **** (p < 0.0001). Scale bars, 10 μm.

We then exploited a FUCCI mouse ESC model ^40^ to discriminate Early G1 (hCdt1++), Late G1/Early S (hCdt1+/Geminin+), Mid/Late S (hCdt1-/Geminin+) and G2 cells (Geminin++) (Fig. 2d). We monitored H3.1/H3.2 patterns in parallel with cell cycle markers. In line with the above results, we observed the lowest H3.1 ‘enriched’ pattern in early G1 and early S phase (∼5% and ∼15% respectively) followed by a strong increase of the same profiles during Mid/Late S and finally G2 (∼80% and ∼95%, respectively) (Fig. 2e). Taken together, these data further support a dynamic equilibrium at chromocenters with H3.1/H3.2 and H3.3 compensating for each other. Furthermore, while differentiated cells show a steady enrichment of H3.1/H3.2 enrichment at chromocenters throughout the cell cycle, ESCs, show a clear loss of H3.1 enrichment during G1 which is only recovered following chromocenter replication.

### DNA synthesis promotes H3.1 enrichment at chromocenters

To investigate key parameters leading to the enrichment of H3.1/H3.2 (but not H3.3) at chromocenters, we considered two aspects. H3 histone variants differ in their amino-acid sequence mainly in two regions: (i) in the histone fold domain: a motif that allows distinct recognition by histone chaperones; (ii) in the amino-terminal tail at position 31: a serine in H3.3 instead of an alanine in H3.1, which phosphorylation proved critical to activate transcription during key cell fate transitions and development ^41–43^. The DNA synthesis coupled (DSC) deposition involves the CAF-1 complex and requires the SVM motif in H3.1/H3.2 ^44–46^. For DNA synthesis independent (DSI) deposition, other histone chaperones interact with the AIG motif in H3.3 ^6,47–49^ (Fig. 3a). Finally, H3.3 also differs from H3.1 by a serine to cysteine substitution at position 96, equally present in H3.2 (Fig. 3a). We generated transgenic ESCs lines with wild-type H3.1, H3.2, and H3.3 SNAP-tag constructs, along with an H3.1 construct with an A31S substitution in its N-terminus (A31S) and an H3.3 with either a phosphomimic (S31D) or phospho-dead (S31A) substitution. We found that all constructs with the SVM motif (corresponding to DSC deposition) showed enrichment at the chromocenters (Fig. 3b). In contrast, constructs with the AIG motif (corresponding to DSI deposition), including the S31 phospho-mimic (S31D) and phospho-dead (S31A) mutants did not accumulate at chromocenters (Fig. 3b, right panels). Our quantification of the proportion of cells displaying ‘enriched’, ‘even’, and ‘excluded’ patterns at chromocenters to S phase progression showed that constructs linked to DSC deposition led to ‘enriched’ patterns (Fig. 3c). The proportion of cells with ‘enriched’ patterns during S phase progression was identical to endogenous H3.1, indicating that the SNAP-HA tag presence did not create any bias (compare Fig. 3c with Fig. 2c and Fig. S1b, ESCs). Importantly, chimera presenting either S31 (from H3.3) or S96 (residues only present in H3.2, Fig. 3a) showed no significant differences; thus, they did not interfere with the stable enrichment. In contrast, the DSI constructs did not show ‘enrichment’ but led to ‘even’ and ‘excluded’ patterns during the S phase like endogenous H3.3 (compare Fig. 3c with Fig. 2c and S1b, ESCs). Importantly, phospho-mimic (S31D) and phospho-dead (S31A) mutants led to similar proportions of ‘even’ and ‘excluded’ patterns compared to their WT counterparts.

**Figure 3.**
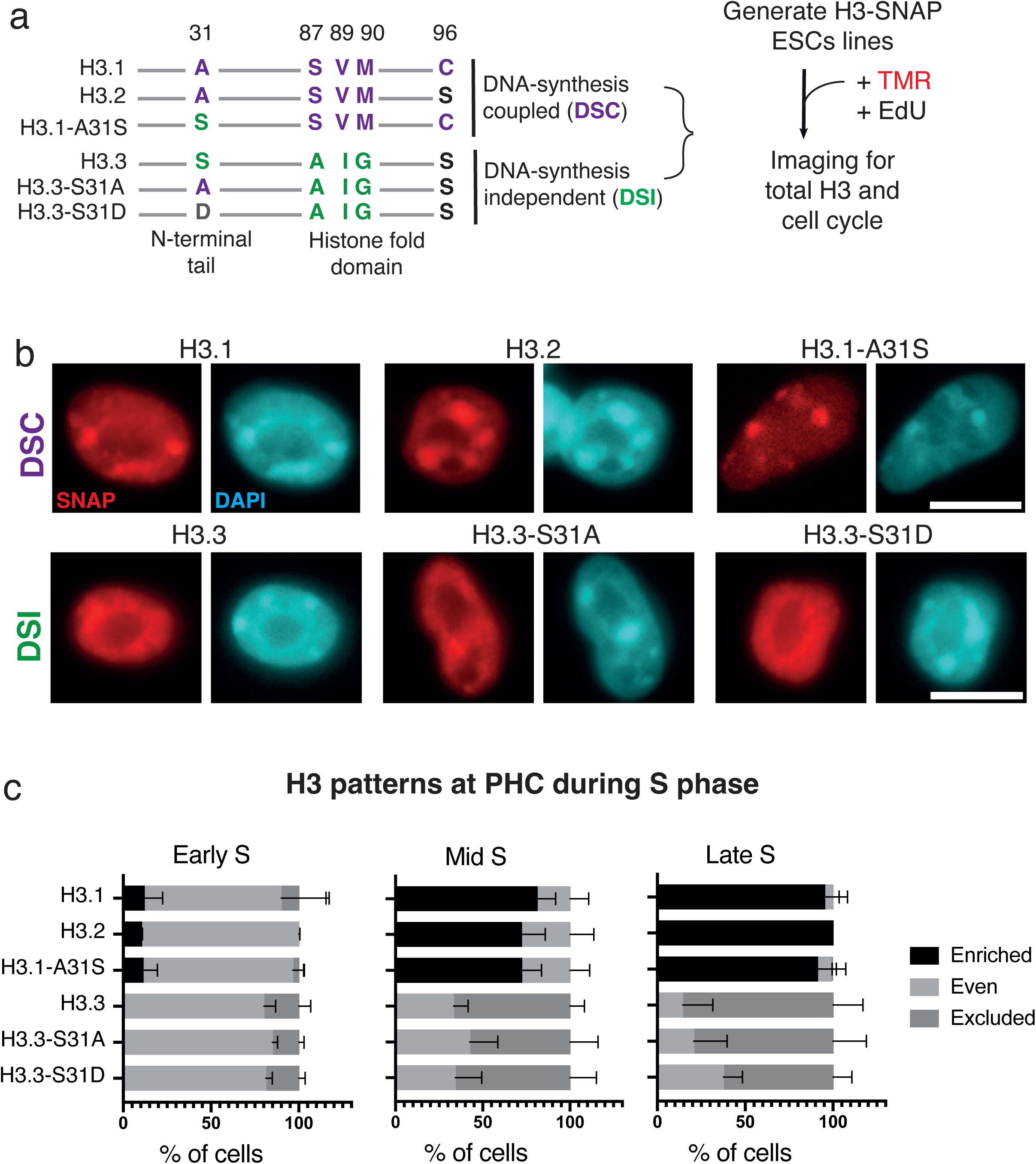
The SVM motif of the HFD is sufficient for H3.1 enrichment at chromocenters. (**a**) Left panel: Specific amino acidic residues are depicted and grouped based on their similarities with the histone chaperone recognition motif. Substitutions from original sequences of H3.1 or H3.3 are highlighted in purple, green, or gray. Right panel: Experimental scheme to obtain ESC lines with point mutations on H3.1/H3.3-SNAP-Tag constructs and visualize total histone H3 patterns upon Pulse (TMR) labeling along with cell cycle. (**b**) Representative epifluorescence images of H3-SNAP-Tag histones (red) in ESCs along with DNA counterstaining (DAPI, cyan). (**c**) Quantification of cells exhibiting H3 patterns at PHC during Early, Mid, and Late S stages. Data show the mean and s.d. (at least 3 biologically independent experiments; >100 nuclei quantified per condition). Scale bars 10 μm.

Several H3 mutants, in cancers are referred to as “oncohistones” ^50^. The amino acid substitutions in the conserved N-terminus of the H3.3 tail, including K27 and G34, are associated with upregulation of neurodevelopmental genes ^51–53^ or mesenchymal differentiation processes ^54–56^. We examined whether the presence of these mutations in H3 variants could affect their subnuclear organization and their enrichment at chromocenters. We thus generated additional ESC lines for H3 oncohistones, including H3.3-K27M, H3.3-K27L, H3.3-G34R, and H3.3-G34V, and monitored the proportion of cells displaying H3 patterns at chromocenters. H3.3 oncohistones like H3.3 did not show any enrichment patterns throughout the S-phase and the highest proportions of even and excluded patterns during Early and Late S, respectively, with little to no differences across the samples (Fig. S3). Notably, they followed the same dynamic changes of the DSI constructs, suggesting that these additional amino acid substitutions did not interfere with the dynamic of H3 depositions at chromocenters. The critical importance of the SVM motif enables us to conclude that the main parameter defining H3.1 enrichment at chromocenters is the DNA synthesis coupled deposition pathway.

### Targeting the HIRA H3.3 chaperone at chromocenters out-competes H3.1 enrichment in ESCs

Since we observed that H3.1 enrichment patterns were mirrored by H3.3 depletion, we reasoned that H3.3 could compete with H3.1 by forcing its deposition at the chromocenters. To test this hypothesis, we engineered a transcription activator-like effector (TALE) designed to bind specifically to the major satellite DNA repeats ^57,58^ in fusion with the H3.3 chaperone HIRA tagged with a Clover fluorescent protein (Fig. 4a). We used both HIRA WT and a mutant isoform (HIRA W700A-D800A) unable to interact with CABIN1 and thus unable to promote *de novo* H3.3 deposition ^59^ (Fig. 4a). We next transfected ESCs and visualized by immunofluorescence endogenous H3.1/H3.2 and H3.3 along with the distinct TALE-HIRA fusion proteins (Clover fluorescence). We first verified that TALE-HIRA and HIRA-W799A-D800A readily localized at the DAPI dense chromocenters as the TALE-Clover (TALE fused to the green fluorescent reporter mClover (TALE-Clover) ^58^, indicating that HIRA fusion proteins did not affect the TALE-mediated specific targeting at chromocenters (Fig. 4a). We next investigated whether the presence of TALE-HIRA impacts H3.1/H3.2 and H3.3 at chromocenters. We observed that cells transfected with TALE-HIRA had a global H3.3 increase in the nucleus (Fig. 4a, right). Most remarkably, we noticed a reduction in the H3.1 signal at chromocenters, which was not the case for TALE-HIRA-W799A-D800A, the mutant deficient for H3.3 deposition (Fig. 4a right). Next, we quantified the proportion of cells displaying ‘enriched’, ‘even’, and ‘excluded’ patterns for H3.3 and H3.1/H3.2 in cells with TALE fusions at chromocenters (Clover+ cells) (Fig. 4b). For H3.3, TALE-HIRA, but not TALE-HIRA-W799A-D800A, increased the proportion of cells with the ‘even’ pattern with a concomitant decrease of cells with the ‘excluded’ pattern (Fig. 4b, left). Remarkably, TALE-HIRA led to a decrease in the proportion of cells displaying H3.1 enrichment at chromocenters (77% to 43%, Fig. 4b right). To confirm this effect, we quantified by 3D-FIED macro the relative enrichment of H3.1/H3.2 and H3.3 at chromocenters targeted by TALE-HIRA and HIRA-W799A-D800A. We analyzed the cells positively enriched for TALE-HIRA or HIRA-W799A-D800A (based on the Clover signal) and used the negatively enriched counterparts as control (Fig. 4c). We found for H3.3 nearly identical enrichment for every condition (Fig. 4d, top panel). We detected a small, but significant increase of H3.3 enrichment when comparing negative and positive cells, for both TALE-HIRA and HIRA-W799A-D800A. However, we did not appreciate any significant difference in H3.3 enrichment between TALE-HIRA and TALE-HIRA-W799A-D800A when we compared positive cells only, indicating that the increased H3.3 enrichment detected with immunofluorescence cannot be appreciated using this quantitative approach (Fig. 4d, top panel). We next performed the same quantification for H3.1/H3.2 and found a significant decrease of H3.1 enrichment specifically in cells positive for TALE-HIRA when comparing 1) negative cells with positive TALE-HIRA cells and 2) positive TALE-HIRA-W799A-D800A cells with positive TALE-HIRA cells (Fig. 4d bottom). Taken together these data indicate that targeting HIRA at chromocenters leads to a small but detectable enrichment of H3.3 at chromocenter (Fig. 4b left) concomitantly with a decrease of H3.3/3.2 (Fig. 4b right, 4d bottom).

**Figure 4.**
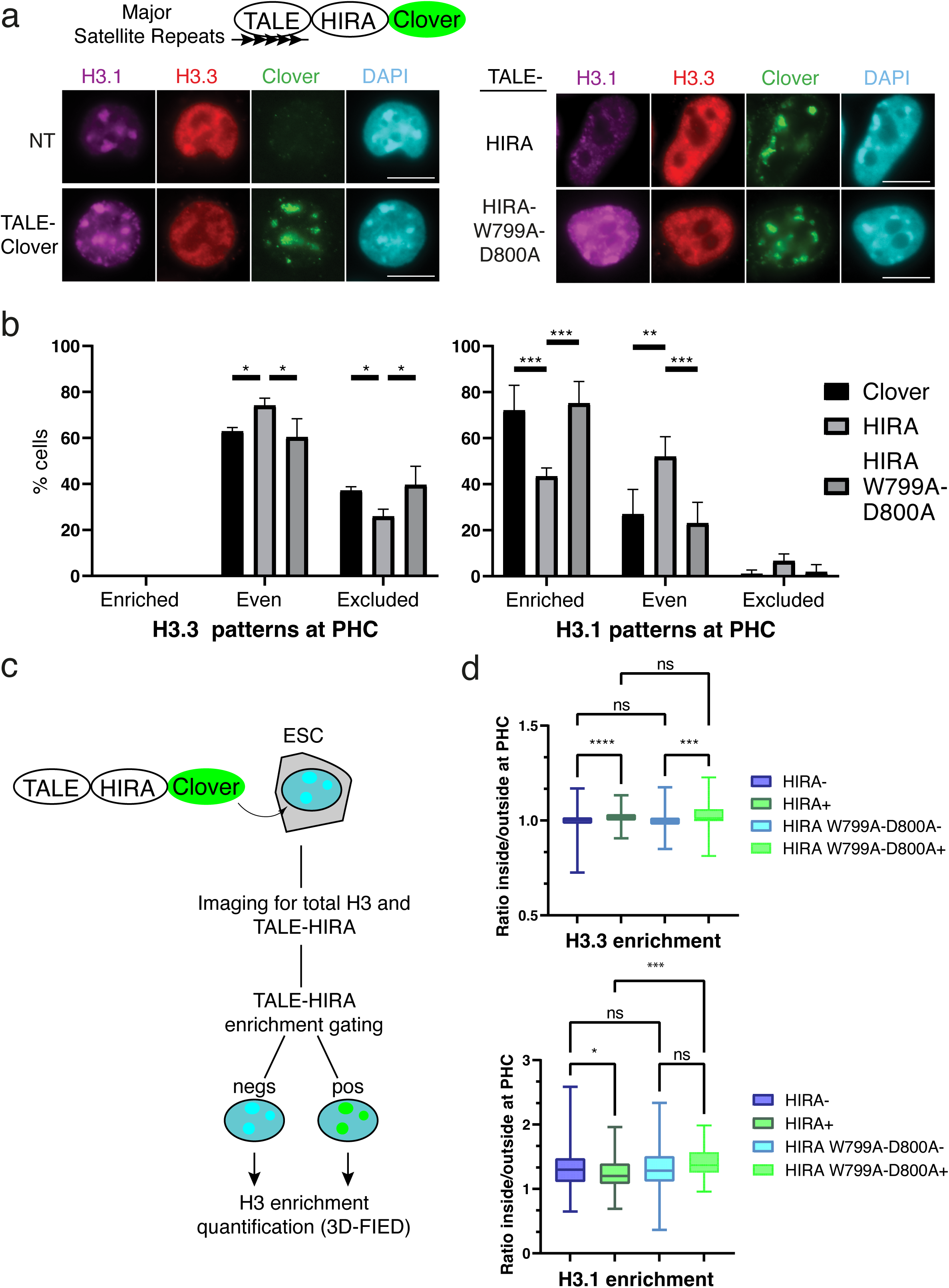
Targeted H3.3 deposition via the HIRA complex alters local H3.1 enrichment at chromocenters. (**a**) Top: Scheme depicting Clover-Tagged HIRA (WT or W799A-D800A) chimeric protein fused to TALE protein recognizing Major Satellite repeat DNA sequences. Left: Representative immunofluorescent images of ESCs transfected with or without TALE-MajSat-Clover control construct. Clover (green) is detected specifically at chromocenters, along with endogenous H3.1/H3.2 (magenta), H3.3 (red), and DNA (DAPI, cyan) staining. Scale bars, 10 μm. Right: same as left, but with TALE-MajSat-HIRA and HIRA-W799A-D800A. (**b**) Quantitative analysis of the percentage of cells exhibiting recurrent patterns of H3 variants at PHC upon transfection of TALE-MajSat -Clover, -HIRA, or -HIRA-W799A-D800A. (**c**) Scheme depicting the experimental approach for gating of TALE-HIRA fusion negative (cyan) and positive (green) cells along with total H3 enrichment quantification by 3D-FIED. (**d**) Boxplots of enrichment values of H3.3 and H3.1/H3.2 at chromocenters in ESCs transfected with TALE-MajSat -HIRA or -HIRA-W799A-D800A. - and +– signs in the legend indicate cells negative and positive cells for TALE-HIRA fusions. Mann-Whitney tests were used for statistical analysis: ns (p > 0.05), * (p < 0.05), *** (p < 0.001), **** (p < 0.0001).

We next wondered if alterations in H3 histone variants balance at chromocenters could impact the centric regions located next to the pericentric regions. We found that the nuclear localization of CENP-A (the histone H3 variant specifically enriched at centric domains) is identical between TALE-Clover, HIRA, and HIRA-W799A-D800A the H3.3 deposition deficient mutant (Fig. S4). This thus indicates that targeting HIRA at chromocenters alters the balance of H3.1/H3.2 and H3.3 at the pericentric regions of the centromere without altering the organization of the centric regions. Taken together, these data support that a targeted HIRA-mediated H3.3 deposition to chromocenters is dominant to out-compete H3.1 presence.

### Targeting the H3.3 chaperone HIRA at chromocenters impacts HP1α, cell division time, and viability

The question that arises is whether impairing H3.1 enrichment at chromocenters affects its function in ESCs and has any impact on cell cycle and mitosis. We thus examined whether targeted H3.3-deposition at chromocenters affects constitutive heterochromatin hallmarks, including HP1α and H3K9me3, and impacts cell viability during cell division. We thus monitored by immunofluorescence H3K9me3 and HP1α at chromocenters in cells targeted by TALE-Clover, -HIRA, and -HIRA-W799A-D800A (Fig. 5a and 5b left). We found enrichment for both H3K9me3 and HP1α at chromocenters in both TALE-HIRA and HIRA-W799A-D800A (Fig. 5a and 5b left). By quantifying enrichment at chromocenters by 3D-FIED, we did not detect significant changes in H3K9me3 enrichment (Fig. 5a right). However, we observed a significant enrichment of HP1α for TALE-HIRA positive cells compared to HIRA negative cells and TALE-HIRA-W799A-D800A cells (Fig. 5b right). While the different behavior between HP1α and H3K9me3 may be surprising, it is interesting to consider that an increase in HP1α may be independent of the modification, possibly linked to other binding properties of HP1α that remain to be explored. Nevertheless, these observations prompted us to consider whether HIRA targeting and H3.1/H3.2 imbalance possibly affect constitutive heterochromatin which could affect its function during mitosis. We tested this hypothesis, by monitoring with time-lapse microscopy the time ESCs spent during cell division and the frequency of cell death events after mitosis. We transfected TALE-Clover, TALE-HIRA, and -HIRA-W799A-D800A in ESCs and followed cell division events for 24h corresponding to at least 1-2 cell divisions (Fig. 5c). We verified that in Clover+ cells the TALE-MajSat fusion proteins were present at chromocenters and pericentric heterochromatin domains during mitosis (Fig. 5c). We determined the cell division time required for cells to progress from late prophase, indicated by a large nucleus with clearly detectable condensed chromosomes, to late telophase, indicated by two well-separated daughter nuclei (Fig. 5c, yellow boxes). In our hands, control cells showed an average cell division time of 45’, whereas for TALE-HIRA transfected cells the average time increased up to 75’ (Fig. 5d). This prolonged time spent in mitosis is accompanied by a 5-fold ratio increase in cell death events compared to control cells (Fig. 5e). Importantly, we did not observe changes in cell division time or death events in TALE-HIRA-W799A-D800A transfected ESCs compared to the control. Thus, the HIRA protein itself does not perturb the DSC-deposition nor cell division process during mitosis.

**Figure 5.**
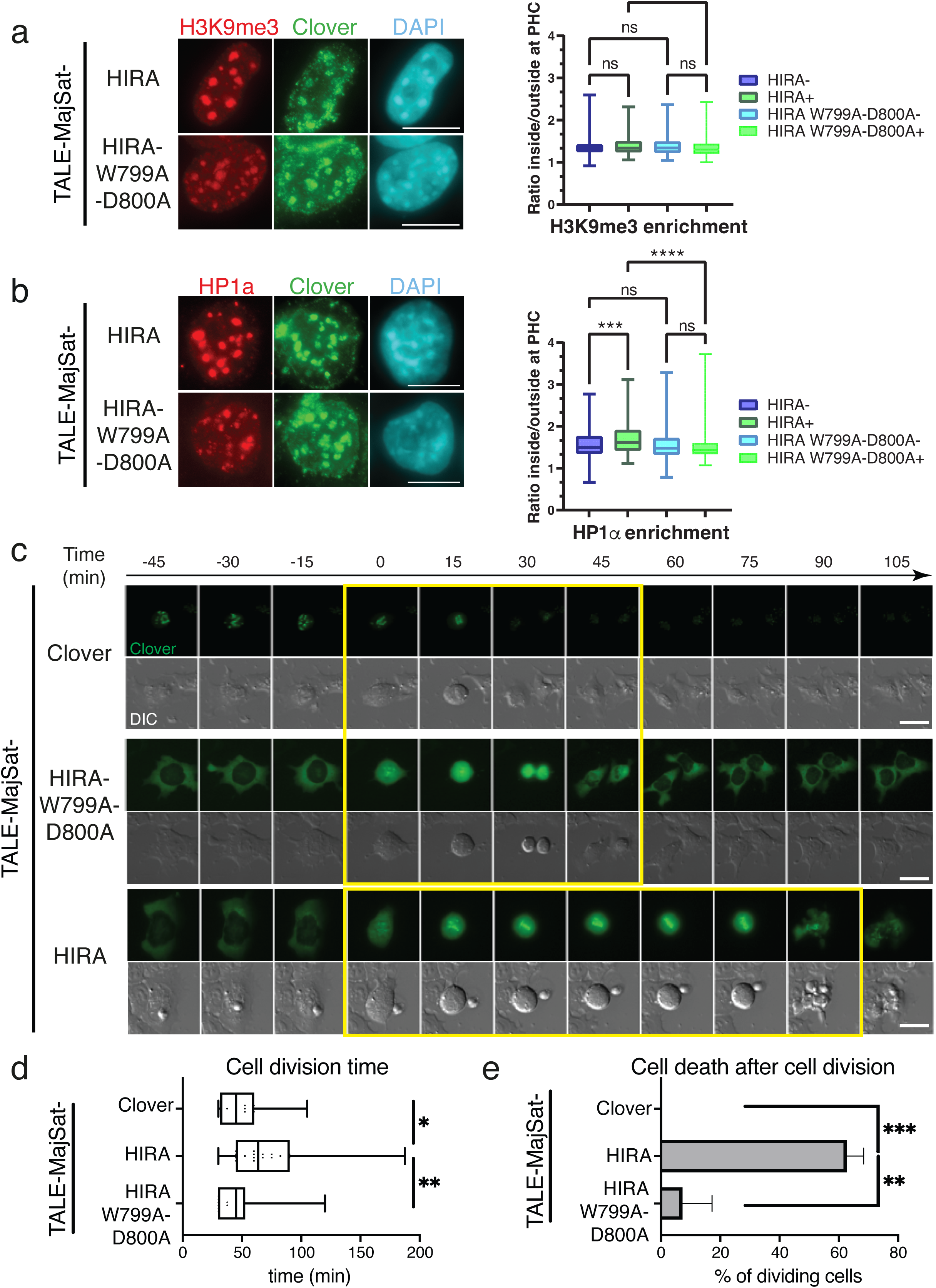
H3.3 deposition at mouse chromocenters impacts cell viability and cell division time. (**a**) Left: Representative immunofluorescent images of ESCs transfected with Clover-tagged TALE-MajSat -HIRA and -HIRA-W799A-D800A constructs. Clover (green) is detected specifically at chromocenters, along with H3K9me3 antibody (red), and DNA (DAPI, cyan) staining. Scale bars, 10 μm. Right: Boxplots of enrichment values of H3K9me3 at chromocenters in ESCs transfected with TALE-MajSat -HIRA or -HIRA-W799A-D800A. - and +– signs in the legend indicate cells negative and positive cells for TALE-HIRA fusions. Mann-Whitney tests were used for statistical analysis: ns (p > 0.05), * (p < 0.05), *** (p < 0.001), **** (p < 0.0001). (**b**) as in (a) but for HP1εξ. (**c**) Series of images showing dividing Clover (green) positive cells of TALE-MajSat-Clover, -HIRA-W799A-D800A, and HIRA by time-lapse microscopy. Differential Interference Contrast (DIC) images boxed in yellow and marked by an arrow are used to determine cell division time in minutes (min). Scale bars, 20 μm. (**d**) Quantitative analysis of the time required to progress from a cell to divide into two daughter cells. The dots represent the mean, and the bars are the min/max values. (**e**) Quantitative analysis of the % of cell death events occurring after one cell division. The bars represent the mean and error bars of the SD. For statistical analysis, we quantified at least 50 cells per condition from 2 biologically independent experiments. Mann-Whiney test: * (p < 0.05), ** (p < 0.01), *** (p < 0.001).

Based on these results, we conclude that the interferences in H3.1 enrichment using a targeted DSI deposition at chromocenters lead to an increase in chromosome segregation defects.

## Discussion

By exploiting a combination of imaging and sequencing methods, we defined how chromocenters show a controlled enrichment of the replicative H3.1/H3.2 rather than the replacement H3.3 histone variant with an important function for cell division and survival. First, we found that chromocenters irrespective of the cell differentiation potential show a reproducible H3.1/H3.2 enrichment paralleled by a relative H3.3 depletion. This enrichment for replicative histone variants depends on the deposition pathway mediated by the CAF-1 histone chaperone in S phase. It is robustly maintained throughout the cell cycle in differentiated cells compared to pluripotent ESCs. In the latter, during the G1 phase, a loss of H3.1 enrichment occurs at the expense of a gain in H3.3 deposition. We experimentally challenged the H3.1 enrichment at chromocenters by forcing H3.3 deposition through the artificial targeting of HIRA to chromocenters (Fig. 6). These conditions compromised heterochromatin marks at chromocenters, and cells showed defects in mitosis and cell cycle. Our data thus indicate a conserved function for the DSC-deposition pathway to sustain H3.1 enrichment at chromocenters and to restrict the DSI-deposition machinery presence at pericentric heterochromatin domains throughout the cell cycle. We discuss the implications of our findings for the specific cell cycles of pluripotent and differentiated cells.

**Figure. 6.**
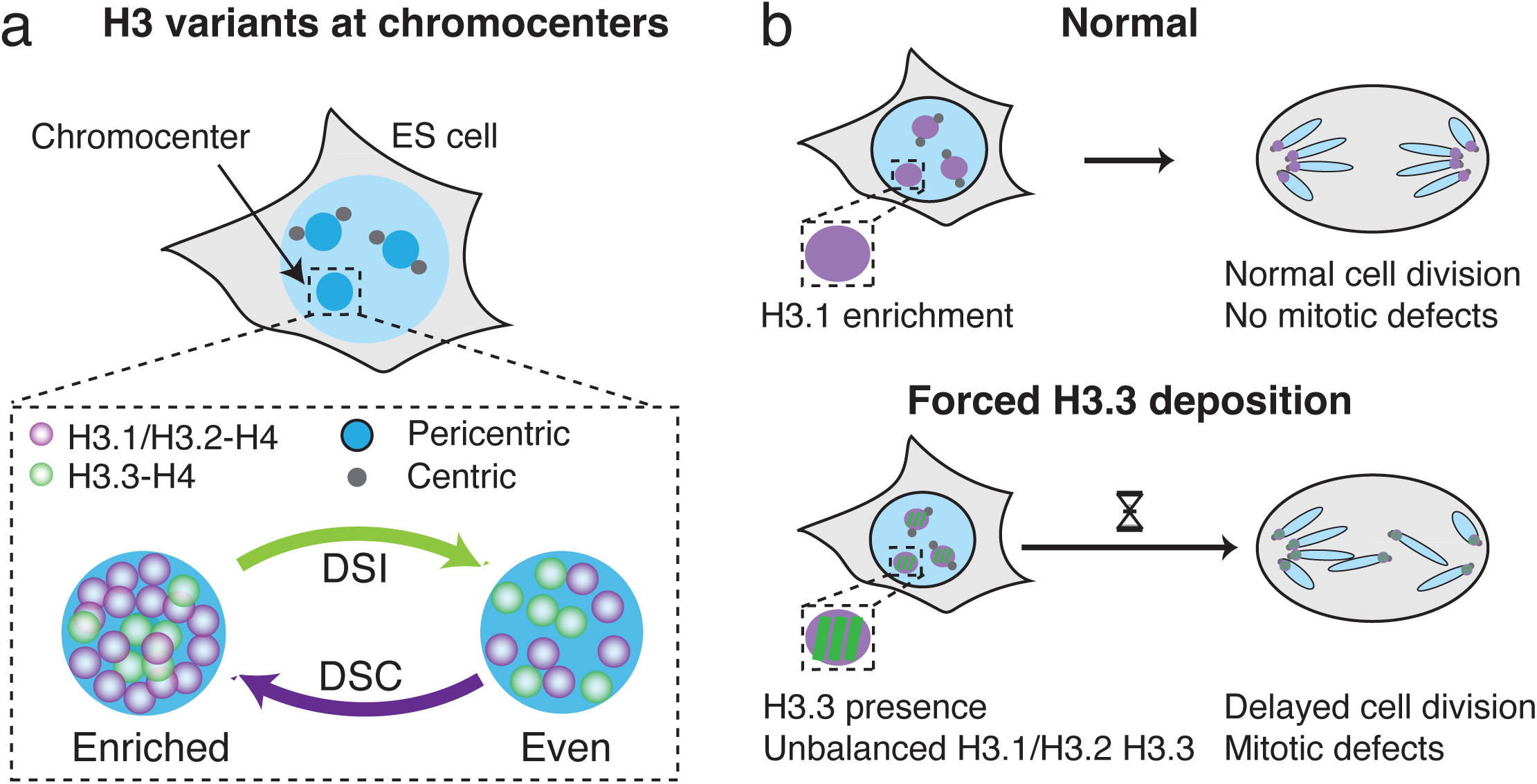
Model. Defects associated with H3.3 targeting via HIRA at chromocenters, in place of H3.1/H3.2 deposition, have severe consequences for ESCs, including chromosome instability and mitotic defects. (**a**): schematic model in interphasic cells of the balance between H3.1/H3.2 and H3.3 at the chromocenter mediated by histone deposition pathways. DSI: DNA synthesis independent (DSI); DSC: DNA-synthesis dependent (DSC) (**b**) Upon unbalanced H3.1/H3.2 and H3.3 at the chromocenter (pericentric regions) mitotic defects ensue.

In mouse cells, the general view posits that replacement variant H3.3 is enriched at active sites, enhancers, and promoters, but also marks heterochromatin close to telomeres and pericentric regions ^6^. However, H3.1 is also present in heterochromatin regions, visibly detected when discernable compartments form in the nuclear space, as shown in our study. Notably, in plants, H3.1 is highly enriched in chromocenters, including the centromeric and pericentromeric chromatin regions ^28,60–62^. These series of observations raise key questions as to how such choices in histone variants with distinct balanced levels can be achieved in nuclear domains. How does the observation in plants relate to mammals? How can the reciprocal dynamics of these variants explain changes associated with cell states during cell cycle and differentiation? We focused on chromocenters to follow the subnuclear distribution of H3.1 and H3.3 in pluripotent ESCs, NPCs, and terminally differentiated cells NIH-3T3 cells. As found in plants ^28^, we detected a consistent H3.1 enrichment at chromocenters and relative H3.3 exclusion. Genome-wide and image analysis confirmed this dual presence of H3.1 and H3.3 at major satellites and close to centromeres as previously reported in human cells and plants ^63–65^. We further dissected by amino acid substitution key residues present in H3.1/H3.2, but not in H3.3, which proved crucial for H3.1 enrichment at chromocenters. This analysis revealed the major importance of the histone fold domain of the SVM motif, which is key for the deposition involving the CAF-1 histone chaperone. Thus, we established that the major mechanism involved in replicative H3 enrichment is strictly dependent on the DNA synthesis deposition pathway. Interestingly, S31, whose phosphorylation is key for the activation of genes during differentiation ^42^, macrophage activation ^41^ and gastrulation in Xenopus ^43^ proved dispensable for this enrichment. Other amino-acid discriminating H3.1 from H3.3 did not prove critical, nor onco-histone mutations.

We then noticed that H3.1 enrichment at chromocenters varied during the cell cycle in a manner mirrored by a reciprocal H3.3 presence (Fig. 2). This relative decrease in the early S-phase reflects the global increase in the H3.1 signal associated with early replication of the euchromatin as opposed to the mid-late replication of heterochromatin in chromocenters (Fig. 2b, left panel, Early S-phase). In agreement with this interpretation, this unbalance is overcome during Mid-S and Late-S when chromocenters replicate and heterochromatin doubles, and this status is maintained in subsequent G2 and propagated in G1. This observation leads us to consider that H3.1 enrichment depends both on its deposition during S phase, but also on the competing deposition of H3.3. For every cell line analyzed, except for ESCs, the early-S phase showed few cells with H3.1 enrichment, and the peak of H3.1 enrichment consistently occurred during the mid-S, late-S, and G2/G1 transition phases. Since euchromatin regions replicate during the early-S phase, increased amounts of H3.1 are expected due to replication-coupled deposition, and this could lead to the relative decrease in H3.1 enrichment at chromocenters which replicate later. Indeed, when chromocenters replicate in the mid-S and late S phase, the relative enrichment between chromocenters and the rest of the nucleus is restored. We found the same behavior for these dynamics of H3.1 relative enrichment in NPCs and NIH-3T3 cells. However, the situation is different in ESCs which transcribe major satellite repeats at higher levels than differentiated cells ^66^. Indeed, preventing major satellite transcription in ESCs enables the formation of stable chromocenters ^67^. Unusual dynamics for replicative H3 are thus observed in ESCs to ensure stable heterochromatin maintenance while preserving pluripotency ability ^26,68,69^. Thus, the dynamics of structural chromatin proteins, including replicative histone variants and HP1, are constantly challenged and need to be replenished in stem cells ^26,27^. Remarkably, when forcing H3.3 deposition we found that chromocenters with less H3.1 acquired higher amounts of HP1α and H3K9me3 in ESCs. This change, with a decrease or a restriction in global HP1α mobility at pericentric domains, has severe consequences during cell division and mitosis (Fig. 6).

While the normal setting of this balance favors replicative histones, it can be challenged and vary according to the cell potential. The question that arises then is whether the more ‘labile’ H3.1 enrichment at chromocenters in ESCs simply reflects a generally higher H3.1/H3.3 turn-over or if it could have a more direct role specifically at chromocenters. Indeed, chromocenters are less well-defined in size and shape in ESCs than in differentiated cells. Interestingly, knocking down the p150 subunit of CAF-1, which impairs H3.1 deposition, leads to a loss of chromocenters in ESCs, but not in differentiated MEF cells ^13^. These contrasting data underline the stronger dependency on the replicative deposition pathway in ESCs, and a stronger maintenance mechanism in place in differentiated cells.

Importantly we found that replicative H3.1/H3.2 and the replacement H3.3 histone variants can both be present at pericentric heterochromatin domains. However, their relative accumulation is a matter of dosage controlled by replicative H3.1 deposition mechanisms as opposed to replacement H3.3 deposition. This histone enrichment at chromocenters depends on the cell type and their cell cycle properties. Therefore, it is tempting to consider that regulating the histone chaperones involved in distinct deposition pathways could represent an attractive means for shaping nucleosomal composition at distinct nuclear domains. Exploring these issues represents exciting avenues to better understand the control in nuclear organization and plasticity during cell fate decisions.

## Materials and Methods

### Cell culture

We cultured all cells at 37°C in 5% CO2: KH2 mESCs ^70^ on gelatinized feeder-free tissue culture plates in ESC media (Dulbecco’s modified Eagle’s medium supplemented with GlutaMax, Pyruvate and 4,5 g/L D-Glucose (Thermo Fisher Scientific), 15% calf fetal serum (Eurobio), 1000 U/ml penicillin/streptomycin, 1X MEM non-essential amino acids, 125 μm beta-mercaptoethanol supplemented with 1000 U/ml LIF (Millipore) and 2i inhibitors, which include 1 μM MEK1/2 inhibitor (PD0325901) and 3 μM GSK3 inhibitor (CHIR99021). To generate NPCs, we cultured mESCs cells in NPC media (ESC media (without LIF and 2i) supplemented with 1 μM Retinoic Acid (RA), 1x N-2 Supplement (Gibco) and 1x B27 Supplement (Gibco) for 4 days, and 3 additional days in the same NPC media supplemented with 10 ng/ml of FGF (Peprotech) and 20 ng/ml of EGF. We cultured NIH-3T3 cells (ATCC #CRL-1658) in Dulbecco’s modified Eagle’s medium (Invitrogen) supplemented with 10% fetal calf serum (Eurobio), 1000 U/ml penicillin/streptomycin (Invitrogen). We cultured FUCCI ESCs (kind gift from D. Landeira) as ^40^.

### Plasmid construction and generation of cell lines

We fused SNAP-3xHA coding sequences downstream H3.1 and H3.3 CDS ^46^ and inserted this fusion protein into the pBS31 vector ^70^ using NEBuilder® HiFi DNA Assembly Master Mix (NEB). These expression vectors (pB31-H3.1- and pB31-H3.3-SNAP-3xHA) enabled the production of H3.1- and H3.3-proteins with the tag SNAP-3xHA at their C-terminus. To generate constructs with point mutations H3.2-, H3.1-A31S- and H3.3-S31A-, H3.3-S31D-SNAP-3xHA plasmids we used site-directed mutagenesis (GenScript) with pB31-H3.1- and pB31-H3.3-SNAP-3xHA vectors, respectively. We integrated the H3-SNAP-3xHA coding sequence in KH2 mouse ESCs downstream of the Type I Collagen (Col1A1) locus containing an Frt site under the control of a TET-ON regulatory region, we co-transfected pB31-H3-SNAP-3xHA plasmids with the pCAGGS-FlpE Vector (Addgene #20733) using Nucleofector Kit 2 (Amaxa) according to manufacturer’s instructions. To generate TALE-HIRA-WT-Clover and TALE-HIRA-W799A-D800A-Clover, we inserted in the pTALYM3B15 plasmid (obtained from Addgene #47878) the HIRA-WT and HIRA-W799A-D800A coding sequences.^59^ using NEBuilder® HiFi DNA Assembly Master Mix (NEB) and verified all constructs by Sanger sequencing. We verified protein expression by Western blot and immunofluorescence analysis. To obtain clones that stably integrated the H3-SNAP-3xHA tag, we selected colonies on hygromycin for 14 days and isolated single clones screened by genotyping and sequencing. We induced histone expression by adding 1 μg/ml doxycycline at least 48h before analysis. We generated NIH-3T3 cells stably expressing either H3.1- or H3.3- SNAP-3xHA as in ^46^ with selection using 10 ug/mL Blasticidin after retroviral transfection.

### Total cell extract preparation and western blotting

We prepared total protein extracts by boiling cell pellet in SDS PAGE loading buffer (Invitrogen) complemented with NuPage reducing agent (Invitrogen) and Universal Nuclease (Pierce) and performed Western blotting as in ^43^ then acquired immunoblot images with ChemiDoc Imager (Biorad). We listed primary antibodies and their preparations in the table (below).

### Immunofluorescence, staining and epifluorescence microscopy acquisition

Cells seeded and grown on fibronectin-coated glass coverslips were transferred into a four-well plate (ThermoFisher Scientific) for labeling. We performed the SNAP-tag labeling *in vivo* with 2 μM SNAP-Cell TMR-Star (New England Biolabs) and visualized histones incorporated into chromatin after a pre-extraction of soluble histones before fixation with 2% paraformaldehyde for 20’ as ^32^. We revealed DNA synthesis by EdU incorporation and Click reaction (Click-iT EdU imaging kit, Invitrogen) as ^32^. Immunofluorescence staining was performed after the Click reaction as ^32^ with the primary antibodies described in the table (below).

### Antibody List and Conditions

| Target | Source of antibody | Specificity | Dilution |
| --- | --- | --- | --- |
| H3.1/3.2 mAb (Clone 1D4F2) | Active Motif, #61629 | Monoclonal Mouse IgG2b | 1:2000 |
| H3.3 | Active Motif, #91191 | Monoclonal Mouse IgG2a | 1:150 |
| HA | Roche, #1867423 | Monoclonal Rat | 1:2000 |
| Aurora B | BD Transduction Laboratories #611082 | Monoclonal Mouse IgG1 | 1:1000 |
| HP1alpha | Euromedex, 2HP-1H5-AS | Monoclonal Mouse | 1:1000 |
| HP1alpha | Sigma, #H2164 | Rabbit Polyclonal | 1:1000 |
| Oct3/4 | BD Biosciences 611203 | Monoclonal Mouse IgG1 | 1:1000 |
| H4 | Abcam ab31830 | Monoclonal Mouse IgG1 | 1:1000 |

Microscopy images, visualization and analysis. We used Fiji software for Microscopy image visualization. To quantify H3.1, H3.3, H3K9me3 and HP1α enrichment at chromocenters, we used the custom 3D-FIED Fiji macro ^33^. We used an enrichment value of 1.2 for TALE- HIRA and HIRA-W799A-D800A (Clover signal) to the gate for the positive (ratio >1.2) and negative cells (ratio <1.2) displaying TALE-HIRA fusion at chromocenters. For quantification of cells with various patterns of H3 variants at chromocenters, we defined cells as (i) ‘enriched’ when they displayed at least three chromocenters for which H3 signal intensity is higher than that of the nucleus; (ii) ‘even distribution when the H3 signal at the chromocenters was that of the nucleus; (iii) ‘excluded’ when the H3 signal at the chromocenters was lower than that of the nucleus. For the quantification of cells within the cell cycle, we identified S phase cells by EdU detection, G2 as EdU negative and AuroraB positive, and G1 as both EdU and AuroraB negative. For quantification of FUCCI ESCs within the cell cycle, we defined Early G1 phase cells as positive for hCdt1 detection, Late G1/Early S positive both hCdt1 and Geminin, Mid/Late S Geminin positive and G2 cells with the strongest Geminin signal. Quantification of cells and fluorescence intensity was performed at least in 100 nuclei for at least three independent experiments unless otherwise indicated. All plots and data visualization are generated with GraphPad and Python. We performed time-lapse microscopy using the Thunder Imaging System (Leica) equipped with a Hit Stage Top Incubation System (Tokai). Briefly, we transfected cells with TALE-MajSat-Clover plasmids and plated on fibronectin-coated dishes (Ibidi) and acquired images every 15 minutes with a 40x objective, during a period of 24h, starting 24h after transfections. We used ImageJ to analyze movies.

### H3.1- and H3.3- SNAP Capture-Seq

We performed H3 ChIP-Seq in ESCs and NPC cells by using the SNAP-capture procedure as in ^34^. We induced synthesis of H3-SNAP-Tag in ESCs and NPCs by adding 1 μg/ml doxycycline before cell collection. We prepared sequencing libraries at the Next Generation Sequencing (NGS) platform at Institut Curie (Illumina TruSeq ChIP kit) and performed PE100 sequencing on Illumina NovaSeq 6000. We used repetitive element annotation from Ensembl (mus musculus core 102_38) and filtered out tandem (‘trf’) and low complexity (‘dust’) repeats classes. For each sample, the number of reads at each repeat was calculated by overlapping the repeat annotation with the genomic coordinates of fragments mapped in pairs, excluding duplicates, extracted from bam files. Samples were normalized to the total number of reads mapped to all repeats (CPM), then by dividing by repeat length, and finally by dividing by the matching input sample. Then, the IP to input ratio was log_2_-transformed, and to allow comparison between conditions, the cross-sample was normalized by computing z-scores.

## Author contributions

G.A., JP.Q. and SA conceived the strategy and wrote the manuscript. G.A and JP.Q. supervised the work. S.A. performed experiments and analyzed data with JP.Q. A.F. and H.M. performed cell biology and chromatin immunoprecipitation experiments for Fig. 1e. T.K. analyzed data in Fig. 1e. E.M. advised on ESC biology. G.A. acquired funding. Critical reading and data discussion involved all authors.

## Acknowledgments

We thank E. Boyarchuk for starting the project, B. Kaffe for generating PBS31_H3.1-SNAP- 3xHA plasmid, and D. Landeira for the FUCCI ESCs. We thank D. Ray-Gallet, and D. Jeffery for critical reading, and all team members and the unit for discussions. We acknowledge the Cell and Tissue Imaging Platform PICT-IBiSA (member of France-Bioimaging ANR-10- INBS-04) in UMR3664 and ICGex NGS platform of the Institut Curie. Funding to G.A. includes La Ligue Nationale contre le Cancer (labellisation), Labex DEEP-PSL (ANR- 11LABX-0044_DEEP, ANR-10-IDEX-0001-02), ERC-2015-ADG694694 ChromADICT and Horizon EIC Pathfinder project 101099654 “RT-SuperES”. Funding to E.M includes Israel Ministry of Science (MOST-DKFZ) collaborative program ([0005358 to EM), Israel Cancer Research Foundation (ICRF) (21-114-PG to EM) and Horizon EIC Pathfinder project 101099654 “RT-SuperES”. S.A. benefited from H2020 MSCA-ITN – EpiSyStem (Grant No. 765966), Fondation Recherche Medicale (FRM) (Grant No. FDT202106012804) and EIC Pathfinder project 101099654 “RT-SuperES”. T. K. benefited from H2020 MSCA-ITN – ChromDesign.

---

**Supplementary Figure 1.**
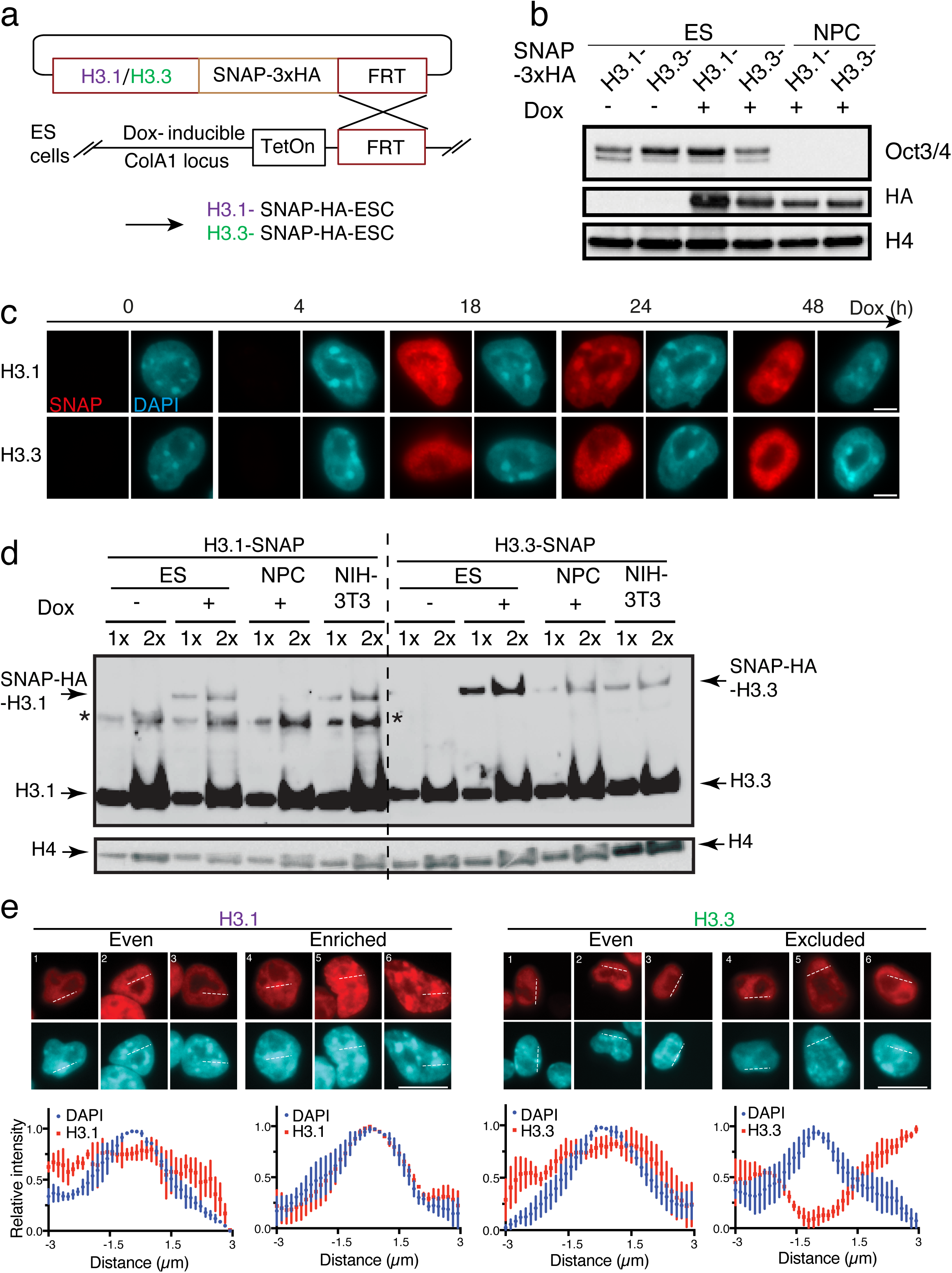
(related to Figure 1**). H3-SNAP-Tagged histones are all expressed and incorporated in a similar mode *in vivo***. (**a**) Schematic and strategy to introduce H3.1 or H3.3 histone variants in mouse ESCs. H3.1 (or H3.3) CDS is fused upstream to a SNAP-Tag-3x-HA sequence and cloned into an expression vector that is stably integrated downstream of the Type I Collagen (Col1A1) locus containing a Frt site under the control of a TET-ON (Dox-inducible) regulatory region. (**b**) Analysis of expression of SNAP-3xHA H3 variants by Western blot of total proteins in ESCs and differentiated NPCs. The presence of SNAP-tagged histones is verified with HA antibody. Differentiation from ESCs to NPCs is verified via Oct3/4 expression. (**c**) Representative epifluorescence images of SNAP-ESCs- H3.1 or H3.3 (red) in the presence (+) or not (-) of Dox along with H3.1/H3.2 or H3.3 staining (magenta) and DNA (DAPI, cyan). (**d**) Comparison by Western blot of endogenous and exogenous expression of H3 histones in ESCs, NPCs, and NIH-3T3 cells. Total protein extracts from 1*10^6 cells (= x) are loaded per lane. The presence of H3 variants is verified by H3.1/H3.2 and H3.3 specific antibodies on two separate membranes at different time exposures. Tagged and endogenous histones are reported at their corresponding heights. H4 histone is used as a loading control. * = non-specific bands. Scale bars 5 μm. (**e**) Representative epifluorescence images of H3.1- (or H3.3-) SNAP (red) ESCs and DNA (DAPI, cyan) with the definition of the three observed patterns (Enriched, Even, Excluded), based on TMR signal intensity in euchromatin and PHC. White dotted lines in red and cyan signal channels define regions of interest to define H3 patterns at chromocenters. Bottom: Dot-scan plots displaying the relative intensities of DAPI and H3.1/H3. IF signal intensity along a 6 μm dashed line traced above a representative pericentric heterochromatin domain.

**Supplementary Figure 2.**
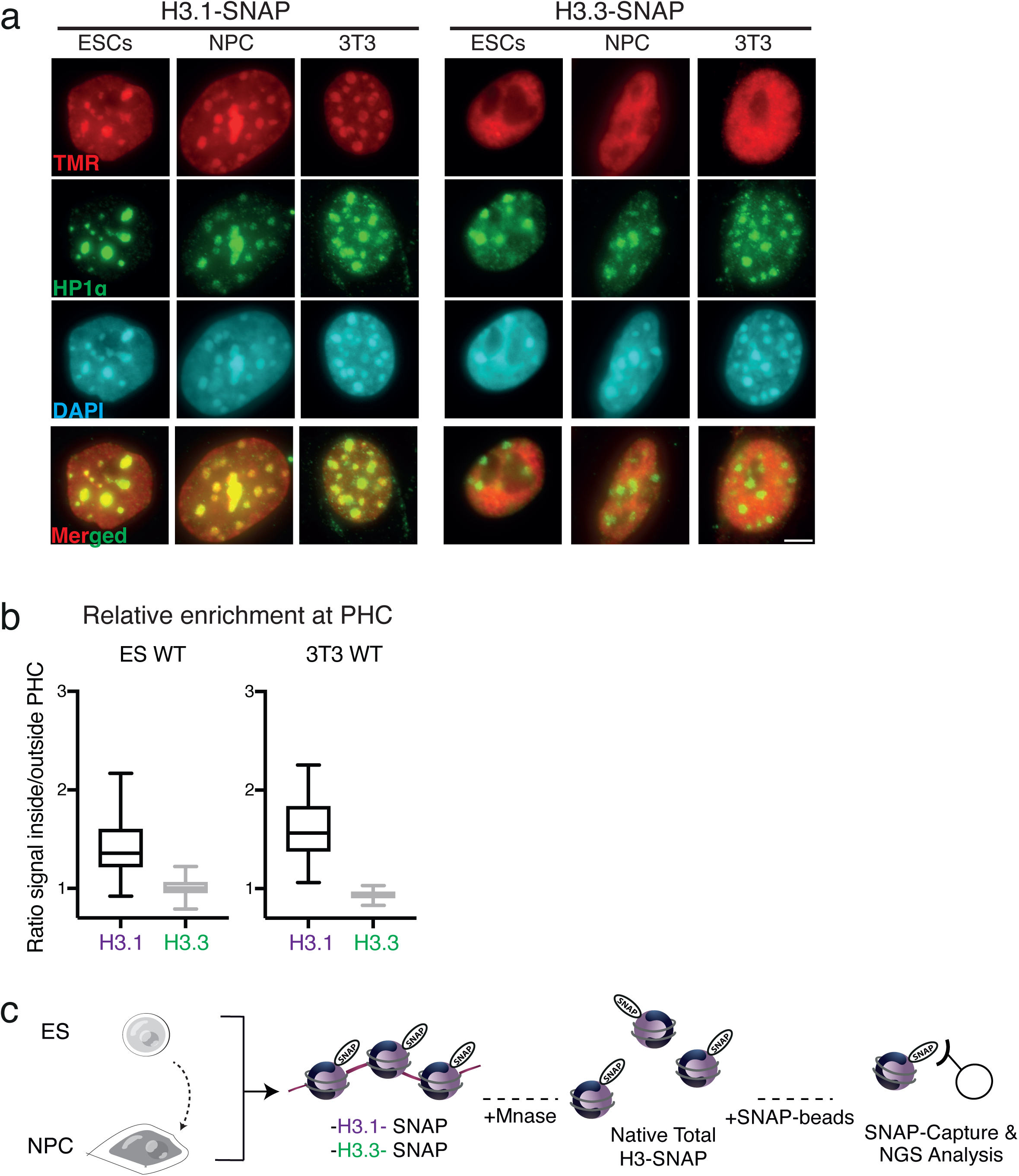
(relative to Figure 1). H3.1 is strongly associated with mouse chromocenters during differentiation. (**a**) Representative epifluorescence images of H3.1- (or H3.3) SNAP (red) in mouse ESCs, NPCs, or NIH-3T3 along with HP1α (green) and DNA (DAPI, cyan). Clusters of pericentric heterochromatin (PHC) were identified as DAPI-dense domains. (**b**) Boxplots of H3.1 and H3.3 immunofluorescence signals with relative enrichment inside of chromocenters in ESCs and NIH-3T3. (**c**) Scheme for SNAP-Seq in mESCs and NPCs expressing H3.1 or H3.3- SNAP. MNase digestion optimized to obtain mononucleosomes next subjected to capture and pull-down with SNAP beads, followed by sequencing.

**Supplementary Figure 3.**
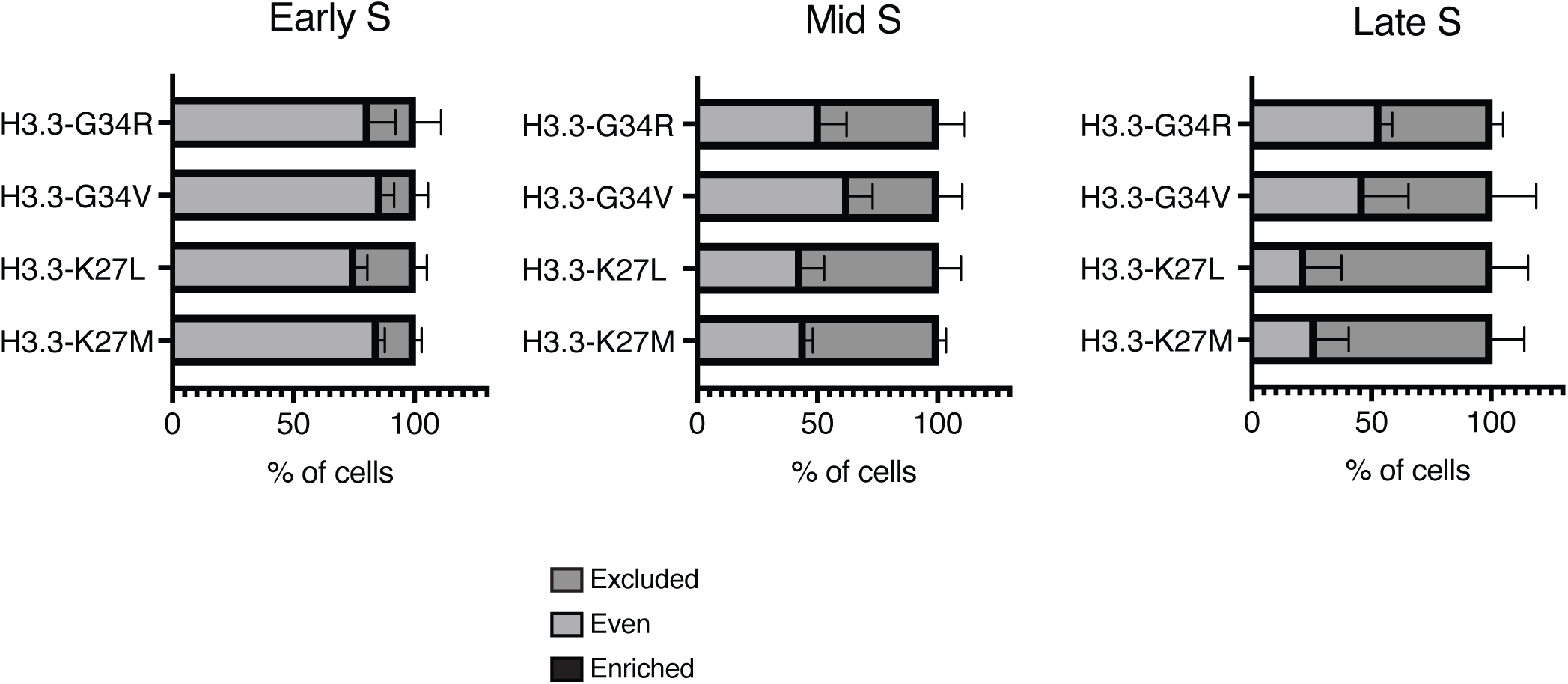
(related to Figure 3**). H3.3-oncohistones are not enriched at chromocenters**. Quantification of cells expressing H3 oncohistones and exhibiting H3 patterns at PHC during Early, Mid, and Late S stages. Data show the mean and s.d. (at least 3 biologically independent experiments; >100 nuclei quantified per condition.

**Supplementary Figure 4.**
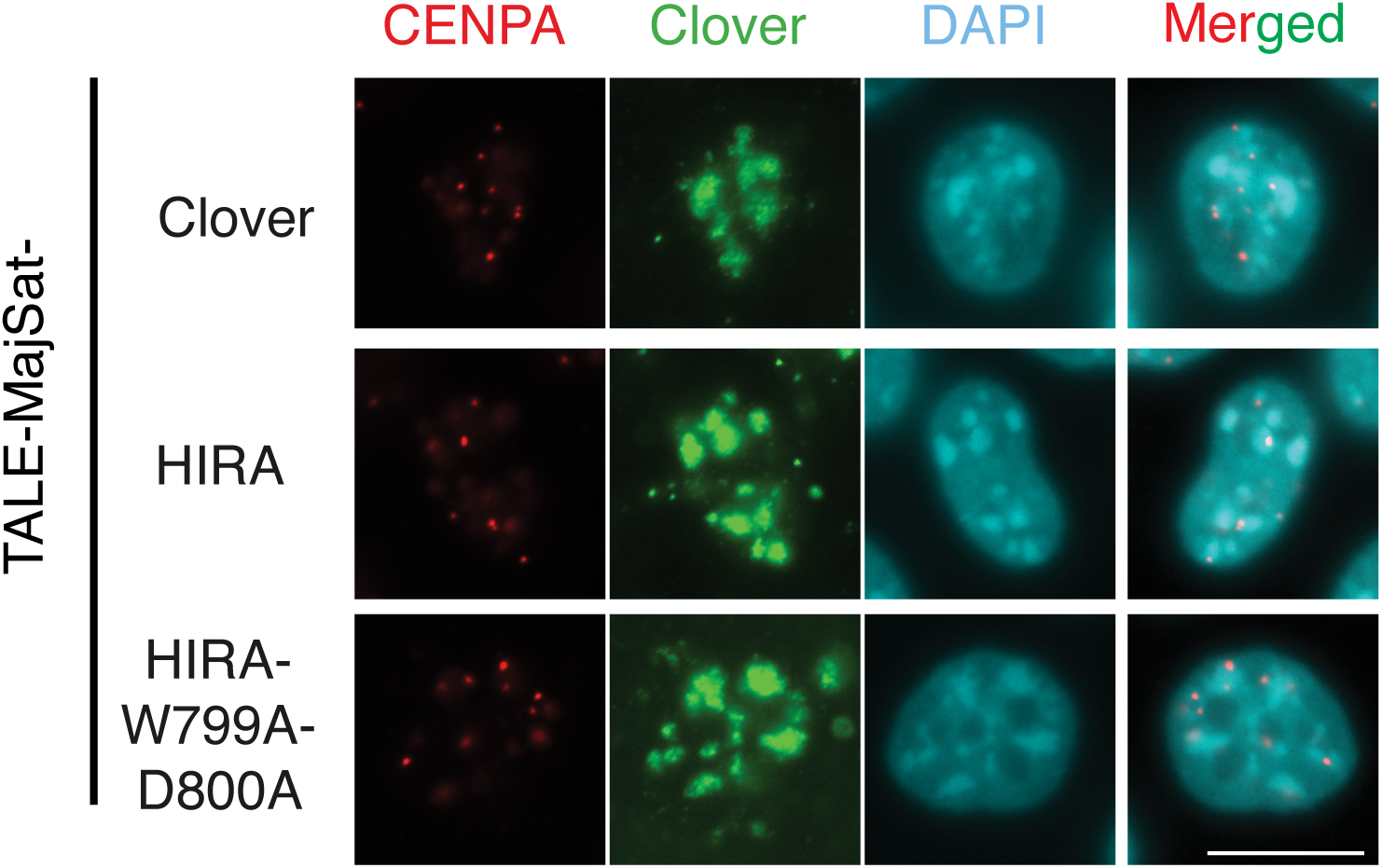
(related to Fig. 4). CENP-A localization in TALE-MajSat-HIRA fusion proteins targeted cells. Representative immunofluorescent images of ESCs transfected with TALE-MajSat-Clover, -HIRA, and -HIRA-W799A-D800A constructs. Clover (green) is detected specifically at chromocenters, along with CENP-A (red) antibody, and DNA (DAPI, cyan) staining. Scale bars, 10 μm.

## References

1. Yadav, T., Quivy, J.-P. & Almouzni, G. Chromatin plasticity: A versatile landscape that underlies cell fate and identity. Science 361, 1332–1336 (2018).

2. Luger, K., Mäder, A.W., Richmond, R.K., Sargent, D.F. & Richmond, T.J. Crystal structure of the nucleosome core particle at 2.8 A resolution. Nature 389, 251–260 (1997).

3. Gurard-Levin, Z.A., Quivy, J.-P. & Almouzni, G. Histone Chaperones: Assisting Histone Traffic and Nucleosome Dynamics. Annual Review of Biochemistry 83, 487–517 (2014).

4. Hammond, C.M., Strømme, C.B., Huang, H., Patel, D.J. & Groth, A. Histone chaperone networks shaping chromatin function. Nature Reviews Molecular Cell Biology 18, 141–158 (2017).

5. Mendiratta, S., Gatto, A. & Almouzni, G. Histone supply: Multitiered regulation ensures chromatin dynamics throughout the cell cycle. The Journal of Cell Biology 218, 39–54 (2019).

6. Goldberg, A.D. et al. Distinct Factors Control Histone Variant H3.3 Localization at Specific Genomic Regions. Cell 140, 678–691 (2010).

7. Elsässer, S.J., Noh, K.-M., Diaz, N., Allis, C.D. & Banaszynski, L.A. Histone H3.3 is required for endogenous retroviral element silencing in embryonic stem cells. Nature 522, 240–244 (2015).

8. He, Q. et al. The Daxx/Atrx Complex Protects Tandem Repetitive Elements during DNA Hypomethylation by Promoting H3K9 Trimethylation. Cell Stem Cell 17, 273–286 (2015).

9. Hoelper, D., Huang, H., Jain, A.Y., Patel, D.J. & Lewis, P.W. Structural and mechanistic insights into ATRX-dependent and -independent functions of the histone chaperone DAXX. Nature Communications 8, 1193 (2017).

10. Drané, P., Ouararhni, K., Depaux, A., Shuaib, M. & Hamiche, A. The death-associated protein DAXX is a novel histone chaperone involved in the replication-independent deposition of H3.3. Genes & Development 24, 1253–1265 (2010).

11. Jagannathan, M., Cummings, R. & Yamashita, Y.M. A conserved function for pericentromeric satellite DNA. eLife 7, e34122 (2018).

12. Peters, A.H. et al. Loss of the Suv39h histone methyltransferases impairs mammalian heterochromatin and genome stability. Cell 107, 323–37 (2001).

13. Houlard, M., Berlivet, S., Probst, A.V. & Quivy, J.-P. CAF-1 Is Essential for Heterochromatin Organization in Pluripotent Embryonic Cells. PLoS Genetics 2, 11 (2006).

14. Maison, C., Quivy, J.-P., Probst, A.V. & Almouzni, G. Heterochromatin at Mouse Pericentromeres A Model for De Novo Heterochromatin Formation and Duplication during Replication. Cold Spring Harbor Symposia on Quantitative Biology 75, 155–165 (2010).

15. Almouzni, G. & Probst, A.V. Heterochromatin maintenance and establishment: Lessons from the mouse pericentromere. Nucleus 2, 332–338 (2011).

16. Burton, A. et al. Heterochromatin establishment during early mammalian development is regulated by pericentromeric RNA and characterized by non-repressive H3K9me3. Nature Cell Biology 22, 767–778 (2020).

17. Casanova, M. et al. Heterochromatin Reorganization during Early Mouse Development Requires a Single-Stranded Noncoding Transcript. Cell Reports 4, 1156–1167 (2013).

18. Probst, et al. A strand-specific burst in transcription of pericentric satellites is required for chromocenter formation and early mouse development. Developmental Cell 19, 625–638 (2010).

19. Probst, A.V., Santos, F., Reik, W., Almouzni, G. & Dean, W. Structural differences in centromeric heterochromatin are spatially reconciled on fertilisation in the mouse zygote. Chromosoma 116, 403–15 (2007).

20. Santenard, A. et al. Heterochromatin formation in the mouse embryo requires critical residues of the histone variant H3.3. Nature Cell Biology 12, 853–862 (2010).

21. Terranova, R., Sauer, S., Merkenschlager, M. & Fisher, A.G. The reorganisation of constitutive heterochromatin in differentiating muscle requires HDAC activity. Experimental Cell Research 310, 344–356 (2005).

22. Solovei, I. et al. Nuclear architecture of rod photoreceptor cells adapts to vision in mammalian evolution. Cell 137, 356–368 (2009).

23. Cohen, L.R.Z. & Meshorer, E. The many faces of H3.3 in regulating chromatin in embryonic stem cells and beyond. Trends Cell Biol (2024).

24. Banaszynski, Laura A. et al. Hira-Dependent Histone H3.3 Deposition Facilitates PRC2 Recruitment at Developmental Loci in ES Cells. Cell 155, 107–120 (2013).

25. Fang, H.-T. et al. Global H3.3 dynamic deposition defines its bimodal role in cell fate transition. Nature Communications 9, 1537 (2018).

26. Meshorer, E. et al. Hyperdynamic Plasticity of Chromatin Proteins in Pluripotent Embryonic Stem Cells. Developmental Cell 10, 105–116 (2006).

27. Boskovic, A. et al. Higher chromatin mobility supports totipotency and precedes pluripotency in vivo. Genes & Development 28, 1042–1047 (2014).

28. Benoit, M. et al. Replication-coupled histone H3.1 deposition determines nucleosome composition and heterochromatin dynamics during Arabidopsis seedling development. New Phytologist 221, 385–398 (2019).

29. Escobar, T.M. et al. Active and Repressed Chromatin Domains Exhibit Distinct Nucleosome Segregation during DNA Replication. Cell 179, 953–963.e11 (2019).

30. Ishiuchi, T. et al. Reprogramming of the histone H3.3 landscape in the early mouse embryo. Nature Structural & Molecular Biology 28, 38–49 (2021).

31. Martini, E., Roche, D.M., Marheineke, K., Verreault, A. & Almouzni, G. Recruitment of phosphorylated chromatin assembly factor 1 to chromatin after UV irradiation of human cells. J Cell Biol 143, 563–75 (1998).

32. Clément, C. et al. High-resolution visualization of H3 variants during replication reveals their controlled recycling. Nature Communications 9, 3181 (2018).

33. Cantaloube, S., Romeo, K., Baccon, P.L., Almouzni, G. & Quivy, J.-P. Characterization of chromatin domains by 3D fluorescence microscopy: An automated methodology for quantitative analysis and nuclei screening. BioEssays 34, 509–517 (2012).

34. Gatto, A., Forest, A., Quivy, J.-P. & Almouzni, G. HIRA-dependent boundaries between H3 variants shape early replication in mammals. Molecular Cell (2022).

35. Forest, A., Quivy, J.P. & Almouzni, G. Mapping histone variant genomic distribution: Exploiting SNAP-tag labeling to follow the dynamics of incorporation of H3 variants. Methods Cell Biol 182, 49–65 (2024).

36. Waisman, A. et al. Cell cycle dynamics of mouse embryonic stem cells in the ground state and during transition to formative pluripotency. Scientific Reports 9, 8051 (2019).

37. Dimitrova, D.S. & Berezney, R. The spatio-temporal organization of DNA replication sites is identical in primary, immortalized and transformed mammalian cells. J Cell Sci 115, 4037–51. (2002).

38. Quivy, J.-P. et al. A CAF-1 dependent pool of HP1 during heterochromatin duplication. The EMBO Journal 23, 3516–3526 (2004).

39. Rausch, C. et al. Developmental differences in genome replication program and origin activation. Nucleic Acids Research 48, 12751–12777 (2020).

40. Asenjo, H.G. et al. Polycomb regulation is coupled to cell cycle transition in pluripotent stem cells. Sci Adv 6, eaay4768 (2020).

41. Armache, A. et al. Histone H3.3 phosphorylation amplifies stimulation-induced transcription. Nature 583, 852–857 (2020).

42. Martire, S. et al. Phosphorylation of histone H3.3 at serine 31 promotes p300 activity and enhancer acetylation. Nature Genetics 51, 941–946 (2019).

43. Sitbon, D., Boyarchuk, E., Dingli, F., Loew, D. & Almouzni, G. Histone variant H3.3 residue S31 is essential for Xenopus gastrulation regardless of the deposition pathway. Nature Communications 11, 1–15 (2020).

44. Ahmad, K. & Henikoff, S. Histone H3 variants specify modes of chromatin assembly. Proc Natl Acad Sci U S A (2002).

45. Latreille, D., Bluy, L., Benkirane, M. & Kiernan, R.E. Identification of histone 3 variant 2 interacting factors. Nucleic Acids Research 42, 3542–3550 (2014).

46. Ray-Gallet, D. et al. Dynamics of Histone H3 Deposition In Vivo Reveal a Nucleosome Gap-Filling Mechanism for H3.3 to Maintain Chromatin Integrity. Molecular Cell 44, 928–941 (2011).

47. Elsässer, S.J. et al. DAXX envelops a histone H3.3–H4 dimer for H3.3-specific recognition. Nature 491, 560–565 (2012).

48. Lewis, P.W., Elsaesser, S.J., Noh, K.-M., Stadler, S.C. & Allis, C.D. Daxx is an H3.3- specific histone chaperone and cooperates with ATRX in replication-independent chromatin assembly at telomeres. Proceedings of the National Academy of Sciences 107, 14075–14080 (2010).

49. Tagami, H., Ray-Gallet, D., Almouzni, G. & Nakatani, Y. Histone H3.1 and H3.3 complexes mediate nucleosome assembly pathways dependent or independent of DNA synthesis. Cell 116, 51–61 (2004).

50. Nacev, B.A. et al. The expanding landscape of ‘oncohistone’ mutations in human cancers. Nature 567, 473–478 (2019).

51. Bender, S. et al. Reduced H3K27me3 and DNA hypomethylation are major drivers of gene expression in K27M mutant pediatric high-grade gliomas. Cancer Cell 24, 660–672 (2013).

52. Lewis, P.W. et al. Inhibition of PRC2 activity by a gain-of-function H3 mutation found in pediatric glioblastoma. *Science (New York*, N.Y.) 340, 857–861 (2013).

53. Mohammad, F. et al. EZH2 is a potential therapeutic target for H3K27M-mutant pediatric gliomas. Nature Medicine 23, 483–492 (2017).

54. Behjati, S. et al. Distinct H3F3A and H3F3B driver mutations define chondroblastoma and giant cell tumor of bone. Nature Genetics 45, 1479–1482 (2013).

55. Bjerke, L. et al. Histone H3.3. mutations drive pediatric glioblastoma through upregulation of MYCN. Cancer Discovery 3, 512–519 (2013).

56. Lu, C. et al. Histone H3K36 mutations promote sarcomagenesis through altered histone methylation landscape. *Science (New York*, N.Y.) 352, 844–849 (2016).

57. Maison, C., Bailly, D., Quivy, J.-P. & Almouzni, G. The methyltransferase Suv39h1 links the SUMO pathway to HP1α marking at pericentric heterochromatin. Nature Communications 7, 12224 (2016).

58. Miyanari, Y., Ziegler-Birling, C. & Torres-Padilla, M.-E. Live visualization of chromatin dynamics with fluorescent TALEs. Nature Structural & Molecular Biology 20, 1321–1324 (2013).

59. Ray-Gallet, D. et al. Functional activity of the H3.3 histone chaperone complex HIRA requires trimerization of the HIRA subunit. Nature Communications 9, 3103 (2018).

60. Stroud, H. et al. Genome-wide analysis of histone H3.1 and H3.3 variants in Arabidopsis thaliana. Proceedings of the National Academy of Sciences 109, 5370–5375 (2012).

61. Vaquero-Sedas, M.I. & Vega-Palas, M.A. Differential association of Arabidopsis telomeres and centromeres with Histone H3 variants. Scientific Reports 3, 1202 (2013).

62. Wollmann, H. et al. Dynamic Deposition of Histone Variant H3.3 Accompanies Developmental Remodeling of the Arabidopsis Transcriptome. PLOS Genetics 8, e1002658 (2012).

63. Dunleavy, E.M., Almouzni, G. & Karpen, G.H. H3.3 is deposited at centromeres in S phase as a placeholder for newly assembled CENP-A in G1 phase. Nucleus 2, 146–157 (2011).

64. Nagaki, K. et al. Chromatin immunoprecipitation reveals that the 180-bp satellite repeat is the key functional DNA element of Arabidopsis thaliana centromeres. Genetics 163, 1221–5 (2003).

65. Shibata, F. & Murata, M. Differential localization of the centromere-specific proteins in the major centromeric satellite of Arabidopsis thaliana. J Cell Sci 117, 2963–70 (2004).

66. Efroni, S. et al. Global Transcription in Pluripotent Embryonic Stem Cells. Cell Stem Cell 2, 437–447 (2008).

67. Novo, C.L. et al. Satellite repeat transcripts modulate heterochromatin condensates and safeguard chromosome stability in mouse embryonic stem cells. Nature Communications 13, 1–16 (2022).

68. Ishiuchi, T. et al. Early embryonic-like cells are induced by downregulating replication-dependent chromatin assembly. Nature Structural & Molecular Biology 22, 662–671 (2015).

69. Nakatani, T. et al. DNA replication fork speed underlies cell fate changes and promotes reprogramming. Nature Genetics (2022).

70. Beard, C., Hochedlinger, K., Plath, K., Wutz, A. & Jaenisch, R. Efficient method to generate single-copy transgenic mice by site-specific integration in embryonic stem cells. genesis 44, 23–28 (2006).

